# Whole-brain studies of spontaneous behavior in head-fixed rats enabled by zero echo time MB-SWIFT fMRI

**DOI:** 10.1101/2021.06.15.448321

**Authors:** Jaakko Paasonen, Petteri Stenroos, Hanne Laakso, Tiina Pirttimäki, Ekaterina Paasonen, Raimo A. Salo, Heikki Tanila, Djaudat Idiyatullin, Michael Garwood, Shalom Michaeli, Silvia Mangia, Olli Gröhn

**Author notes:** Corresponding author. A.I.V. Institute for Molecular Sciences, University of Eastern Finland, P.O. Box 1627, FI-70211, Kuopio, Finland.

## Abstract

Understanding the link between the brain activity and behavior is a key challenge in modern neuroscience. Behavioral neuroscience, however, lacks tools to record whole-brain activity in complex behavioral settings. Here we demonstrate that a novel Multi-Band SWeep Imaging with Fourier Transformation (MB-SWIFT) functional magnetic resonance imaging (fMRI) approach enables whole-brain studies in spontaneously behaving head-fixed rats. First, we show anatomically relevant functional parcellation. Second, we show sensory, motor, exploration, and stress-related brain activity in relevant networks during corresponding spontaneous behavior. Third, we show odor-induced activation of olfactory system with high correlation between the fMRI and behavioral responses. We conclude that the applied methodology enables novel behavioral study designs in rodents focusing on tasks, cognition, emotions, physical exercise, and social interaction. Importantly, novel zero echo time and large bandwidth approaches, such as MB-SWIFT, can be applied for human behavioral studies, allowing more freedom as body movement is dramatically less restricting factor.

## Introduction

Understanding the link between the (ab)normal brain activity and behavior, during, e.g., motivation^1^, emotions^1,2^, or decision-making^3^, is one of the key challenges of modern neuroscience. Rodent studies are traditionally considered well-suited for this purpose, as rodents exhibit many basic aspects of mammal behavior and cognitive processing, such as detection, discrimination, and categorization of sensory information^4,5^. Additionally, rodent studies allow controlled and advanced study designs including surgical, pharmacological, and genetic manipulations.

The relationship between brain activity and behavior has typically been studied with invasive electrophysiological recordings in head-restrained or freely moving rodents^4^. Positron emission tomography (PET)^6,7^, optical imaging^4^, functional ultrasound (fUS)^8^, and functional magnetic resonance imaging (fMRI)^9^ have also been exploited. While electrophysiological and optical measurements provide high temporal resolution, they suffer from limited spatial coverage. PET provides whole-brain coverage but suffers from low temporal resolution. fUS allows high temporal and spatial resolution, but only partial brain coverage has been demonstrated^10,11^ and the limited bone penetration restricts its use in larger species. Awake rodent fMRI provides good spatial and temporal resolution with whole-brain coverage^12–14^, but the subject is typically fully restrained to minimize head and body movement-induced artefacts, making complex behavioral study designs not achievable. Thus, behavioral studies currently lack tools to provide insights into how the whole brain network is modulated during complex behavior.

While studying behavior with fMRI, body movement restriction and loud acoustic noise are the greatest confounders in both animal and human applications. To overcome these and several other limitations imposed by the standard commonly utilized echo planar imaging (EPI) fMRI, we introduced a novel 3D zero echo time fMRI approach based on the Multi-Band SWeep Imaging with Fourier Transformation (MB-SWIFT) technique^15–17^. MB-SWIFT fMRI does not rely on the blood oxygenation level dependent (BOLD) contrast, but appears to be sensitive to blood inflow^16^ and provides even better functional contrast as compared to spin-echo EPI during deep brain and spinal cord stimulation^16,18^. Additionally, we demonstrated that MB-SWIFT signal correlates with neuronal activity^17^. Importantly, we showed that MB-SWIFT is in principle ideal for behavioral studies, as it is up to 20 dB quieter compared to EPI due to small gradient steps, and it is insensitive to magnetic field inhomogeneities due to large bandwidth in all three directions, making it highly immune to movement- and susceptibility-induced artefacts that can severely confound EPI-based fMRI in experimental settings with body movement or surgical preparations^16,17^.

In the present work, we demonstrate that MB-SWIFT fMRI enables unprecedented whole-brain behavioral studies in spontaneously behaving head-fixed rats. We show detailed and anatomically relevant functional parcellation covering both cerebrum and hindbrain, and sensory, limb movement, exploration, and stress-related brain activity in relevant networks during corresponding behavior at both group and individual level. We propose that a transition from the use of traditional EPI sequences to novel zero echo time and large bandwidth sequences can not only significantly improve the fMRI data quality by minimizing scanner noise, motion artefacts, and image distortions, but also open new fMRI study designs including behavioral responses and social interaction in animals and in humans.

## Results

### Habituation of Head-Fixed Rats for fMRI

Sprague-Dawley rats were selected for the experiments because of their capability to adapt quickly for head-fixation with minimal experienced stress^19^. For head-fixation, anchoring screws and layers of bone and dental cement were applied on the skull, leaving two pin holes penetrating the implant horizontally (Figure 1A). The space between the pin holes was large enough for positioning a chronic electroencephalography (EEG) connector (Figure 1B) for simultaneous EEG-fMRI, and can be used together with many other concurrent stimulation or recording techniques.

**Figure 1.**
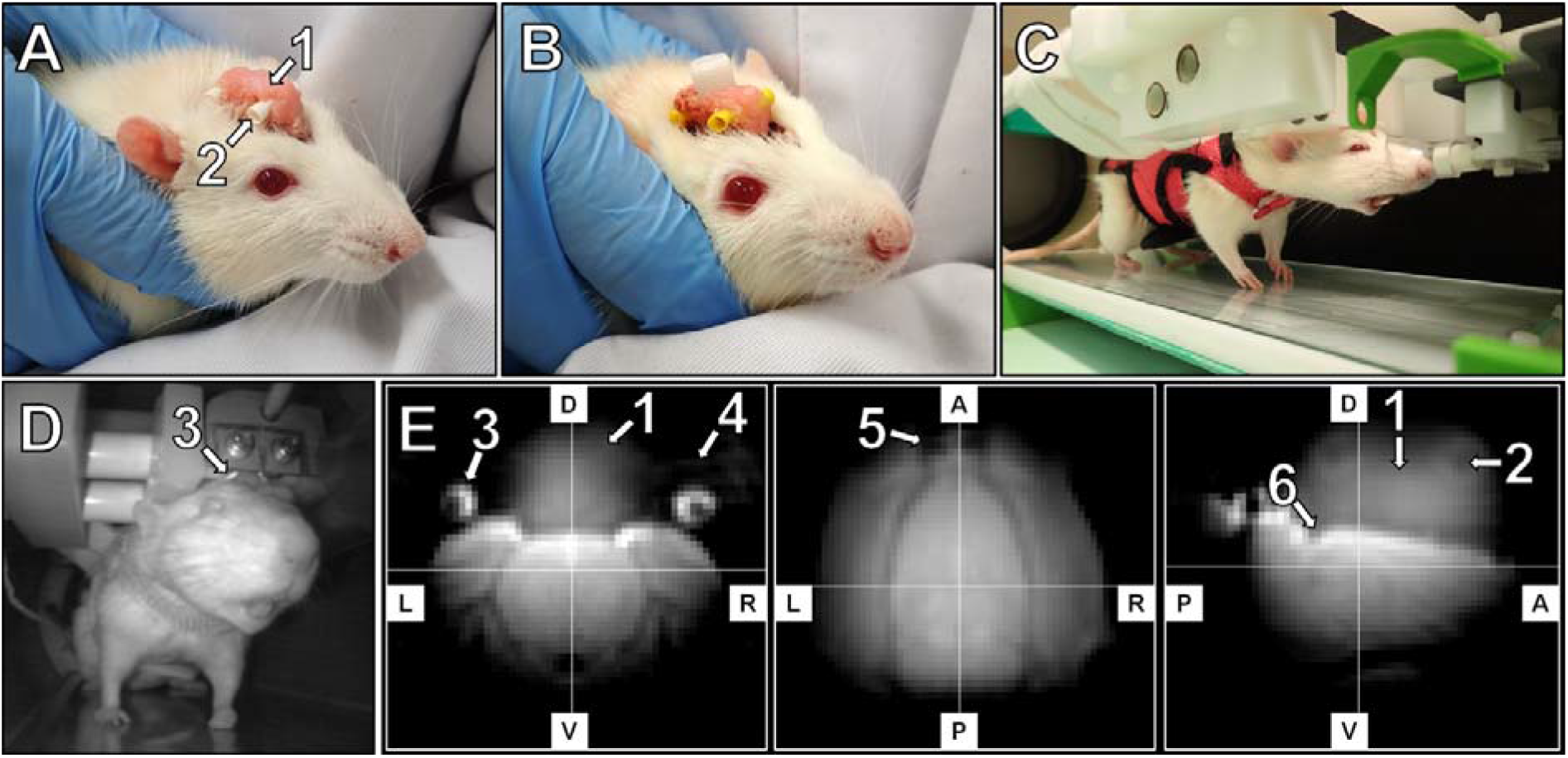
The head implant (A) with an optional electroencephalography electrode connector (B), a head-fixed anesthetized rat wearing a walking harness in the habituation and imaging holder (C), a habituated rat inside the 9.4T MRI scanner (D), and representative MB-SWIFT fMRI images with 625 μm isotropic resolution obtained from an awake head-fixed rat (E). The implant (1) and two 2-mm pin holes (2) for head-fixation were visible in the MB-SWIFT images (E). The custom-made single-loop transmit-receive radiofrequency coil (3) was placed under the two 2-mm pins (4) fixing the head. Anchoring screws on top of both olfactory bulb (5) and cerebellum (6) provided additional support for the bone and dental cement (1) adhesion. Warm-water circulation kept the platform of the holder warm (C). A, anterior; D, dorsal; L, left; P, posterior; R, right; V, ventral.

To prevent excessive pressure on the head implant, rats wore a walking harness that allowed, e.g., standing, sitting, and body and limb movement, but maintained the body alignment (Figure 1C). A video camera was placed to follow behavior, and a custom-made radiofrequency coil was fitted around the implant (Figure 1D). Resulting unprocessed MB-SWIFT fMRI images of head-fixed awake rat show full brain coverage with minimal to no distortions originating from the anchoring screws or interfaces between air, implant, and/or skull (Figure 1E), indicating excellent image quality.

To minimize stress and movement during the head-fixed fMRI, the rats were gradually habituated for imaging in a mock scanner according to the protocol described in Supplementary Figure 1A. We found that corticosterone levels increased (1.75 ± 0.35-fold; p = 0.001, paired two-tailed t-test) and weight decreased (2.8 ± 1.3 %; p < 0.001, paired two-tailed t-test) during the habituation (Supplementary Figure 1B and 1C). However, these suggest that rats were experiencing only mild stress, as awake rodent fMRI studies report up to 6-fold^20,21^ and stress studies up to 10-fold^22,23^ peak increases in corticosterone levels, and restraint-induced body weight loss can typically be 10 %^22^. We observed no difference in the corticosterone levels between the last measurement during the habituation period and the 1^st^ fMRI (p = 0.45; paired two-tailed t-test), suggesting that the habituation environment mimicked well the imaging environment.

The mild stress level of the rats was also supported by the breathing rate (122 ± 23 breaths per minute (bpm); averaged across all data, n = 21 and 15-25 min each), which corresponded to normal non-stressed awake state^24^ and was stable throughout the fMRI measurements (Supplementary Figure 1D). The fMRI was started when either breathing rate exceeded 100 bpm or rat moved spontaneously. At this point the cortical EEG signal had transitioned from initial anesthetized state, required for the imaging preparations, to theta-oscillations (4-8 Hz) typical for awake state (Supplementary Figure 1E). Taken together, the data suggest that the rats were fully conscious during data acquisition.

Additionally, we measured sound pressure levels during both MB-SWIFT and traditional EPI fMRI sequence, and found that MB-SWIFT had 16.0 dB lower (p < 10^-12^, two-tailed t-test) average sound pressure level in the current setup, which is well in line with our previous findings^17^ and beneficial in reducing stress and discomfort during fMRI.

### Spontaneous Behavior and Head Movement

While in the magnet, the rats performed various spontaneous activity, e.g., body twitches, limb movement, sniffing, whisking, eye-related activity, or teeth grinding. Our controlled body movement experiments suggest that small body position changes in head-fixed rats can significantly affect B_0_ field and subsequently EPI-based fMRI images, while raw MB-SWIFT images show minimal movement-induced signal changes in the brain (Supplementary Figure 2). Importantly, the distortions in EPI images were difficult or impossible to correct with commonly applied post-processing methods.

To assess the MB-SWIFT data quality in spontaneously behaving head-fixed rats, we analyzed the motion correction parameters and the corrected data (Supplementary Figure 3). When the translation and rotation parameters between adjacent volumes were analyzed (n = 21 datasets), we found negligible volume-wise average movement (7.8 ± 2.0 μm) and rotation (0.04 ± 0.01°) of the head. Additionally, we found that both the maximum head displacement (0.46 ± 0.32 voxels corresponding to 280 ± 200 μm) and maximum rotation (1.33 ± 1.28°) remained low across datasets. These suggest good head-fixation, as average head motion of 20-30 μm and maximum head displacement of 200-1000 μm was previously reported in head-fixed restrained rats^21^. Nevertheless, the small changes in head position in our study likely arise from the use of slightly flexible MRI-compatible materials (plastic and fiberglass) in the custom-made head-fixation system.

In addition, we studied the temporal stability of head position (Supplementary Figure 3). We observed that 87.8 ± 14.2% of volumes in each data (n = 21) were translated less than 0.2 voxels (125 μm) from the original position, and 91.6 ± 14.7% of volumes were rotated less than 0.5° from the original direction, also indicating good general stability of the head-fixation throughout the measurements. Manual inspection after motion correction led to exclusion of only 3.0 ± 2.4% of volumes because of blurring or remaining visible motion. This provided the vast majority of the data for functional analyses, suggesting excellent overall data quality.

Lastly, we found out that simple rigid transformation was sufficient for motion correction purposes and scaling factors were not required. This minimized potential scaling-induced artefacts during motion correction but also supported the concept that MB-SWIFT images are not affected by the body movement-induced field distortions in contrast to traditional EPI, where severe body movement-induced artefacts can remain despite affine motion correction and regression of correction parameters (Supplementary Figure 2).

### Functional Parcellation and Clustering

To evaluate the general functional data quality, we investigated whether a data-driven approach produces anatomically and functionally relevant brain parcellation in the head-fixed rats. A group-level independent component analysis (ICA) including all task-free data (n = 21, 450-750 volumes each) was performed, and the result (Figure 2 and Supplementary Figure 4) was compared to a rat cerebrum functional parcellation obtained from a larger cohort of awake restrained animals^25^. Despite our smaller data and simpler analysis approach, we found strikingly similar localization of parcels (Figure 2B) to the study by Ma et al. (2018). Compared with the described categorization^25^, we observed similar components in olfactory, prefrontal, cingulate, motor, striatal, somatosensory, auditory, visual, thalamic and hypothalamic, hippocampal and retrohippocampal, midbrain, and brainstem regions. There were only minor differences, as we did not detect a component covering the amygdala, but in contrast observed components covering olfactory bulb (Figure 2B: 1, 4, and 5), insula (Figure 2B: 4), and inferior colliculus (Figure 2B: 31). Overall, the results indicate high functional data quality obtained from the head-fixed behaving rats and robust reproducibility of the awake rat cerebrum functional parcellation across laboratories.

**Figure 2.**
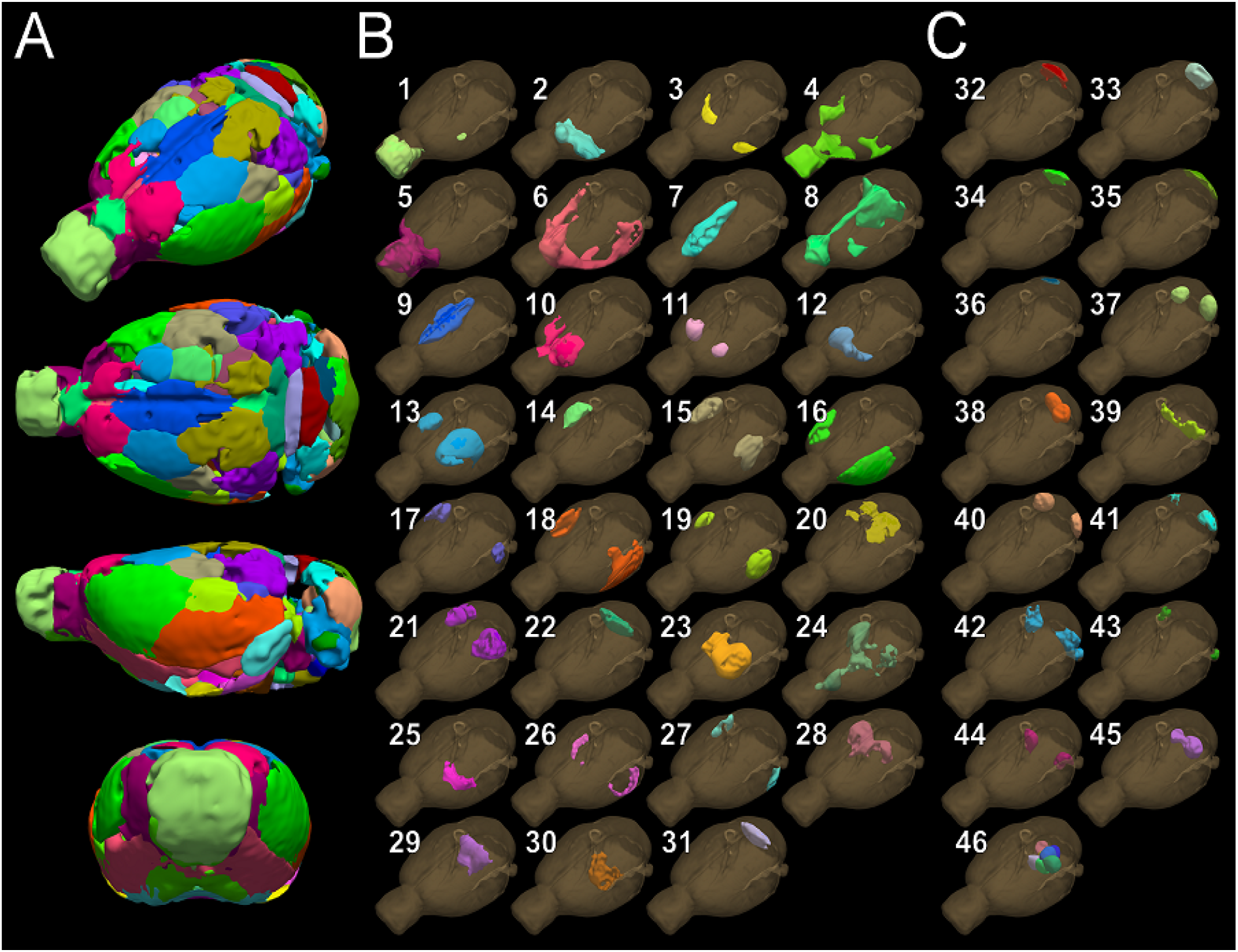
Independent component analysis (ICA) based functional parcellation (A) of cerebrum and olfactory bulb (B), and hindbrain (C), obtained from awake head-fixed rats. Components (B) were in good agreement with the previously described functional parcellation of the rat brain (Ma et al., 2018), which was categorized as follows: olfactory (1-5), prefrontal (6-7), cingulate (7, 9), motor (10), striatal (11-12), somatosensory (13-17), auditory (18-19), visual (20-22), thalamic and hypothalamic (23-25), hippocampal and retrohippocampal (26-28), midbrain (28-29), and brainstem (29-30) areas. Component covering inferior colliculus (31) was also observed. In addition, ICA suggested functional parcellation of the hindbrain (C) as follows: cerebellar lobules (32-39), Crus1 and Crus2 (40-41), flocculus and paraflocculus (42-43), trigeminal nuclei (44), and other brainstem nuclei (45-46). Analysis included all task-free data (n = 21, 15-25 min or 450-750 volumes each). ICA was performed separately for cerebrum (40 components) and hindbrain (30 components), and clear motion-related components were excluded. Unilateral components having a clear contralateral pair were combined to minimize the number of images, and the Z-score threshold given by the ICA was adjusted at least two times stricter for more focused visualization of the components. Subfigure 46 combines several small components in the brainstem. Individual components are shown in Supplementary Figures 4 and 5 in more detail.

The previous work has, however, focused only on rat cerebrum and excluded hindbrain, likely due to the limited coverage of the traditional 2D EPI sequence, low signal-to-noise ratio, and/or image distortions. The cerebellum plays a key role in fine tuning and coordination of movements and its role in cognitive functions is increasingly emphasized^26^. To fill this gap, we show here the first functional parcellation of awake rat hindbrain, consisting of cerebellar lobules, Crus1 and Crus2, flocculus and paraflocculus, trigeminal nuclei, and smaller brainstem nuclei (Figure 2C, Supplementary Figure 5). These observations suggest that the functional subregions of rat hindbrain may also be reliably mapped in behaving rats with MB-SWIFT fMRI, enabling novel functional whole-brain study designs.

To study the functional connectivity across the parcels of head-fixed rat brain, we used the obtained ICA components (Figure 2) as data-driven regions-of-interest (ROIs) and calculated partial correlation coefficients between their fMRI time series. Because 9 rats underwent the task-free imaging session twice, we first tested whether the obtained functional connectivity matrices were reproducible across sessions. We found no differences (p > 0.30, paired two-tailed t-test with multiple comparison correction and Fisher Z-transformation) between the connectivity matrices obtained during the first and second imaging session (Supplementary Figure 6). This observation allowed us to pool the data together for subsequent connectivity analyses.

To study the connectivity structure of head-fixed rats, an average correlation matrix was first calculated from whole data. Subsequently, ROI pairs were organized based on their correlationvalues by using hierarchical clustering (Figure 3A), and the correlation matrix was re-organized accordingly (Figure 3B). As a result, we observed 8 distinct modules (Figure 3C) as follows: 1) hypothalamus and ventral hippocampus, 2) cochlear nuclei and posterior temporal association cortex, 3) posterior hypothalamus, brainstem and cerebellum, 4) medial entorhinal cortex and septal hippocampus, 5) posterior midline cortex, tectum, and dorsal cerebellum, 6) anterior cingulate and somatosensory cortices, 7) septum, thalamus, and cingulate cortex (with sagittal sinus), and 8) olfactory system, prefrontal cortex, and motor cortex. Based on prior anatomical and functional information, components in modules 1, 4, 6, 7, and 8 belong partially or fully into relevant functional systems. For example, in module 4 the hippocampal-entorhinal^27^ and in module 8 the olfactory^28^ and salience^29^ systems were detected. The clustering result shares similarities with a previous report in restrained rats^30^. However, modules 2, 3, and 5 consist mainly of small regions that are anatomically in close vicinity. These findings indicate that large and behaviorally relevant functional networks can be detected and classified robustly in head-fixed rats, but the current resolution may not be sufficient to functionally segregate small regions in deep structures. Additionally, module 7 includes a component that overlaps both sagittal sinus and cingulate cortex, which may also arise from the limited image resolution.

**Figure 3.**
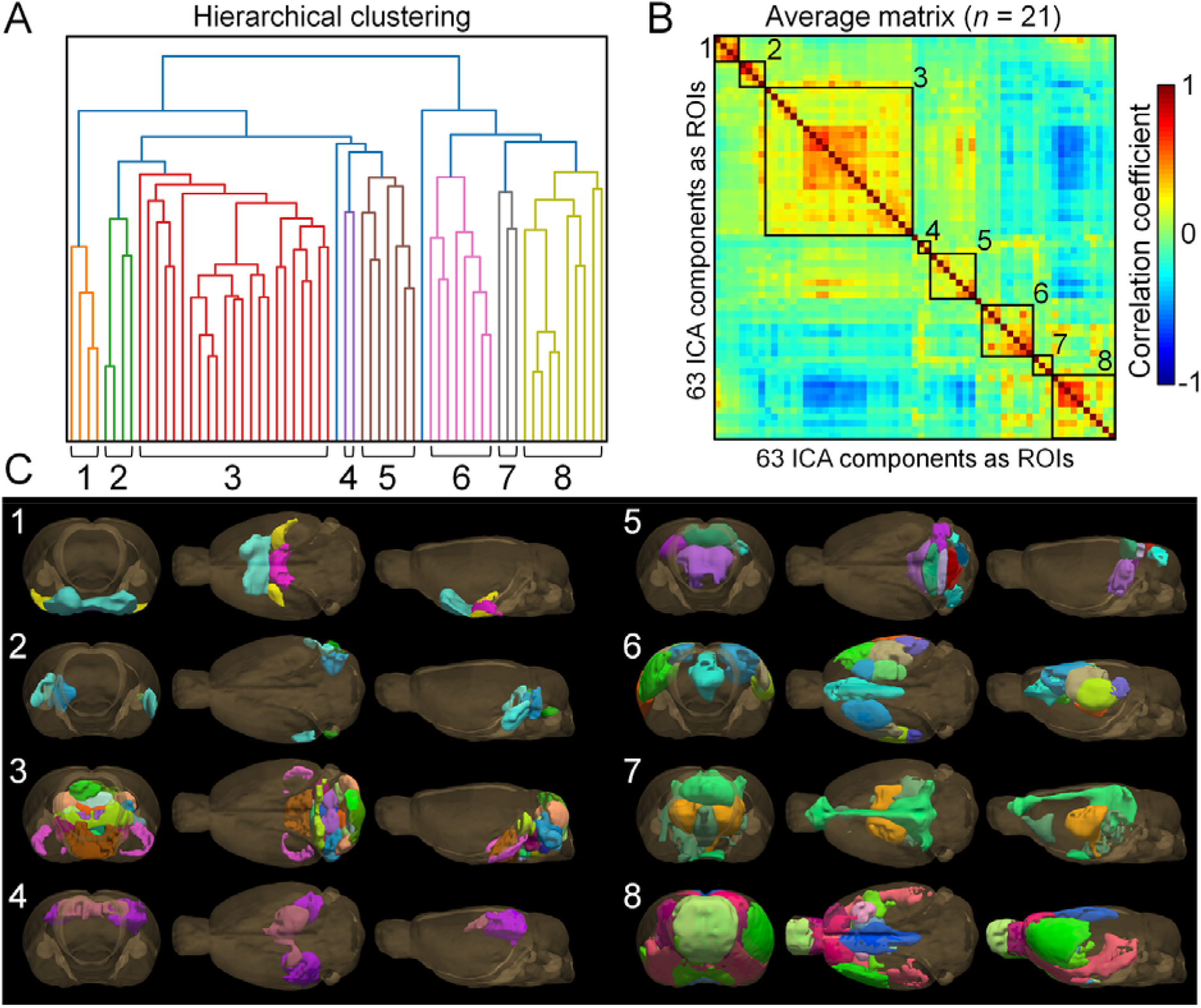
Hierarchical clustering of the correlation data into modules (A), clustered average correlation matrix obtained from task-free fMRI data with modules highlighted (B; n=21), and illustration of functional modules of behaving head-fixed rats (C). Regions-of-interest (ROIs) were derived from the data-driven independent component analysis. Subsequently, partial correlations across ROI pairs were calculated using motion regression parameters and global signal as regressors. Lastly, hierarchical clustering was applied to divide the regions into functional modules, resulting in 1) hypothalamus and ventral hippocampus, 2) cochlear nuclei and posterior temporal association cortex, 3) posterior hypothalamus, brainstem and cerebellum, 4) medial entorhinal cortex and septal hippocampus, 5) posterior midline cortex, tectum, and dorsal cerebellum, 6) anterior cingulate and somatosensory cortices, 7) septum, thalamus, cingulate cortex, and sagittal sinus, and 8) olfactory system, frontal cortex, and motor cortex. The 3D illustrations show views from the front, top, and left side of the brain, respectively. ICA, independent component analysis; ROI, region of interest.

### Group-Level Analysis of Spontaneous and Odor-Induced Behavior

During the fMRI, the rats were mostly passive, although exhibited spontaneous activity. To study the whole-brain activation patterns during spontaneous behavioral events, we selected the three most commonly occurring types of events that were similar: retreat attempt (n = 19), teeth grinding or bruxing (n = 331), and sniffing (n = 19). The onset and duration of the event was obtained from the simultaneously recorded video and used in the event-based group-level fMRI analysis. Examples of these behavior categories are shown in Supplementary Video 1.

Results of the group-level event-based fMRI analyses are shown in Figure 4 (A-C). We found that when the rats suddenly tried to move away from the imaging holder, fMRI signal changes were observed in motor areas, such as motor cortex, cerebellum, and ventromedial and ventrolateral thalamic nuclei^31^ (Figure 4A). Additionally, signal changes occurred in sensory limb regions, as expected, but also in regions involved in decision-making and stress, such as orbitofrontal cortex^32^, cingulate cortex^33^, and posterior hypothalamus^34^, potentially driving the initiation of movement and subsequent motor output.

**Figure 4.**
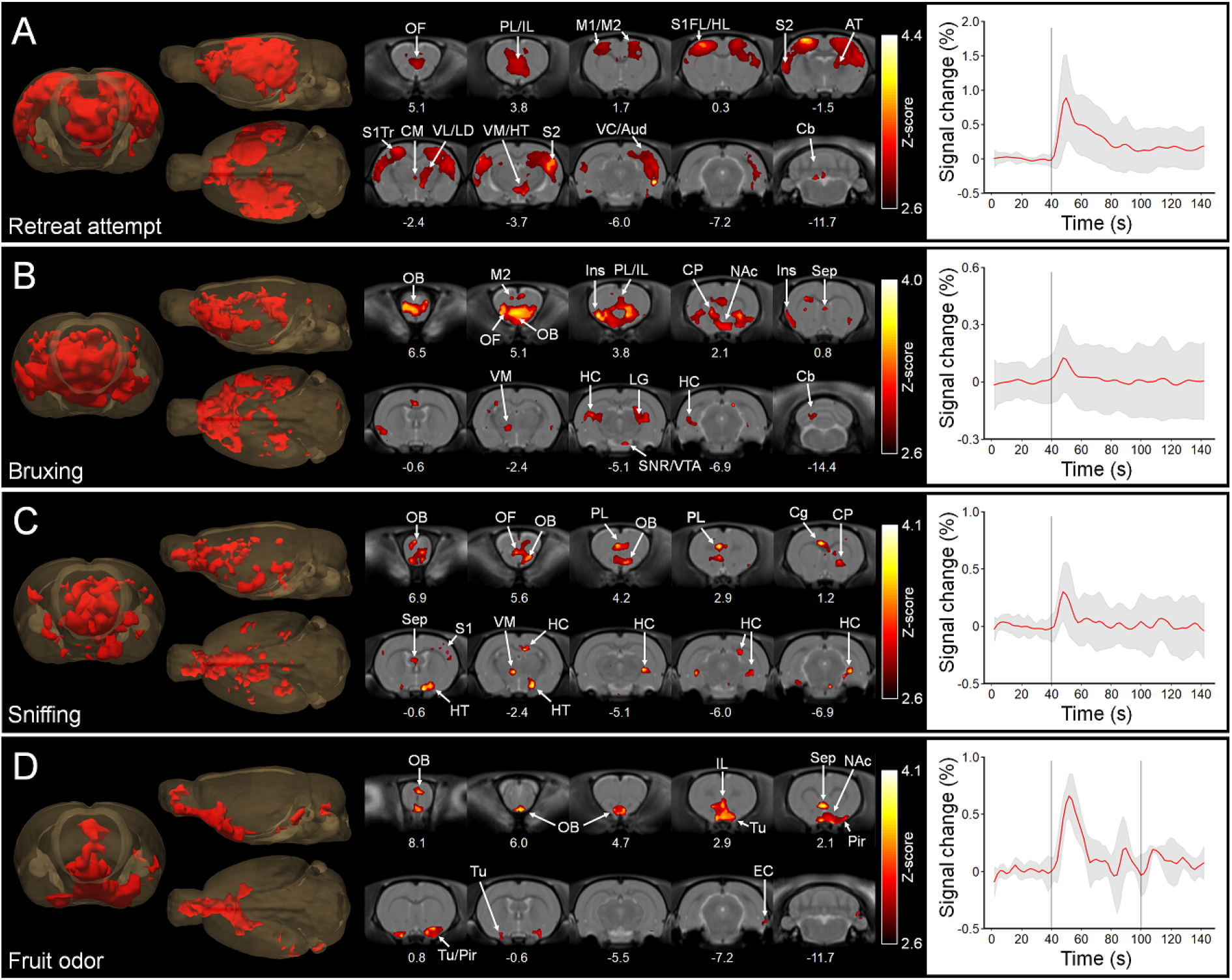
Group-level statistical maps (p < 0.005, corrected) indicating the behavior-related localization of fMRI signal changes (A-C), and the fMRI response to fruit odor (amyl acetate) challenge (D). In A, the rat tried to retreat from the imaging holder (n = 19), leading to signal changes in cortico-thalamic somatomotor areas and the medial frontal cortex. In B, the jaw movement was linked with grinding of teeth (bruxing, n = 331), with simultaneous signal changes in basal ganglia and anxiety-related brain regions (cingulate, insula, and orbitofrontal cortex). In C, the rat was sniffing spontaneously (n = 19), inducing signal changes in olfactory bulb but not in piriform cortex. In D, the rat was sniffing as it was exposed to fruit odor through the nose cone (n = 8), leading to widespread signal changes in the olfactory system including piriform cortex. The grey bars in the time series (average ± standard deviation across all significant voxels) indicate the onset of the observed behavior in the simultaneously recorded video (A-C), and in D the start and end of the odor stimulus. Statistical maps, obtained with FSL FEAT, are overlaid on SIGMA rat brain template. Time series are obtained from events that produced significantly activated voxels. Numbers below slices indicate distance from bregma (mm). The 3D illustrations show views from the front, left side, and top of the brain. AT, anterior thalamic nuclei; Aud, auditory cortex; Cb, cerebellum; Cg, cingulate cortex; CM, centromedial thalamus; CP, caudate putamen; EC, entorhinal cortex; FL, forelimb region; HC, hippocampus; HL, hindlimb region; HT, hypothalamus; IL, infralimbic cortex; Ins, insular cortex; LD, laterodorsal thalamic nuclei; LG, lateral geniculate nucleus; M1, primary motor cortex; M2, secondary motor cortex; NAc, nucleus accumbens; OB, olfactory bulb; OF, orbitofrontal cortex; Pir, piriform cortex; PL, prelimbic cortex; S1, primary somatosensory cortex; S2, secondary somatosensory cortex; Sep, septum; SN, substantia nigra; Tr, trunk region; Tu, olfactory tubercle; VC, visual cortex; VL, ventrolateral thalamus; VM ventromedial thalamus; VTA ventral tegmental area.

The typical purpose of bruxing is teeth maintenance, but it can also be a sign of stress^35^. When the rats were bruxing during the fMRI, we observed localized activation in extrapyramidal motor areas, such as caudate putamen, ventromedial thalamic nuclei, cerebellum, and substantia nigra (Figure 4B), likely related to the typical motor activity during bruxing. Moreover, we observed signal changes in brain regions related to avoidance reactions and anxiety such as insula, anterior cingulate cortex, nucleus accumbens, (lateral) septum and ventral hippocampus^36^. These observations suggest that the cause for bruxing here was mild stress likely induced by the imaging environment. We also observed partial activation of the olfactory system (olfactory bulb and anterior olfactory area). As olfactory (piriform) cortex was not activated and there was no visible sniffing, we hypothesize that the activation in frontal olfactory areas may arise from the altered air flow in nose during the bruxing rather than because of active sniffing.

Sniffing is one of the fundamental exploratory actions in rodents^28,37^. As expected, the olfactory bulb was activated during spontaneous sniffing (Figure 4C). However, the olfactory tubercle and piriform cortex were not activated, suggesting minor odor sampling in the absence of delivered odor in a stable laboratory environment. Instead, the activated regions included the medial frontal cortex, caudate putamen, and hippocampus, which are more related to exploratory behavior in general.

For further investigation, the rats were given gaseous amyl acetate, referred as a fruit odor, through the nose cone (n = 8). In contrast to exploratory spontaneous sniffing, fruit odor-induced sniffing elicited a clear response in the olfactory tubercle, piriform cortex, and lateral entorhinal cortex (Figure 4D), indicating active odor sampling. The fMRI response decayed relatively fast after the onset, likely reflecting the fast olfactory adaptation^38^. The behavioral response to the fruit odor varied between mild and active sniffing across the rats, and we found high correlation between the fruit odor-induced behavioral response and fMRI response strength (r = 0.92, p = 0.0012; Supplementary Figure 7). This further indicated robust coupling between the observed behavior and measured fMRI signal.

Importantly, when the data were reanalyzed without using a brain mask, we found that there were minimal signal changes outside the brain (Supplementary Figure 8), indicating robustness of the findings and minimal confounding effect of movement on the results.

### Individual Spontaneous Behavioral Events

Because of the high individual-level variability in the expression of behavior, there were many other distinct event types that either occurred only rarely or were not considered similar enough to be pooled together for group-level analyses. As such data are anticipated in behavioral studies, we investigated whether the novel approach has sufficient sensitivity to detect relevant brain activity during single behavioral events. In case of borderline statistical significance, we applied two thresholds during the analyses, where one was with the standard (p < 0.05, corrected) and the other with slightly lower (p < 0.001, uncorrected) significance threshold.

To evaluate the sensitivity of the method for single behavioral events, we first chose three whisking-related events, as whisking is another important exploratory action in rodents^37^. The first event is shown in Supplementary Video 2. We found that in all individual-level cases the activation of key regions^39^, such as barrel cortex, motor cortex, and ventral posterior thalamus, was detected during whisking (Figure 5, A-C). The activation of other regions in the tactile somatosensory pathway^39^, such as secondary somatosensory cortex (Figure 5, A and C) and trigeminal nuclei (Figure 5A), were also detected with higher inter-individual variability. In addition to barrel circuit, we observed simultaneous sniffing-related activation of olfactory areas (Figure 5B) or eye movement-related activity in auditory cortex and cerebellum (Figure 5C), suggesting that the eye movement may have been elicited by a sound. Finally, we observed signal changes in regions related to orientation towards stimuli (inferior and superior colliculus^40,41^). These findings indicate consistency between the exploratory behavior and fMRI signal changes observed in multiple networks during whisking-related activity.

**Figure 5.**
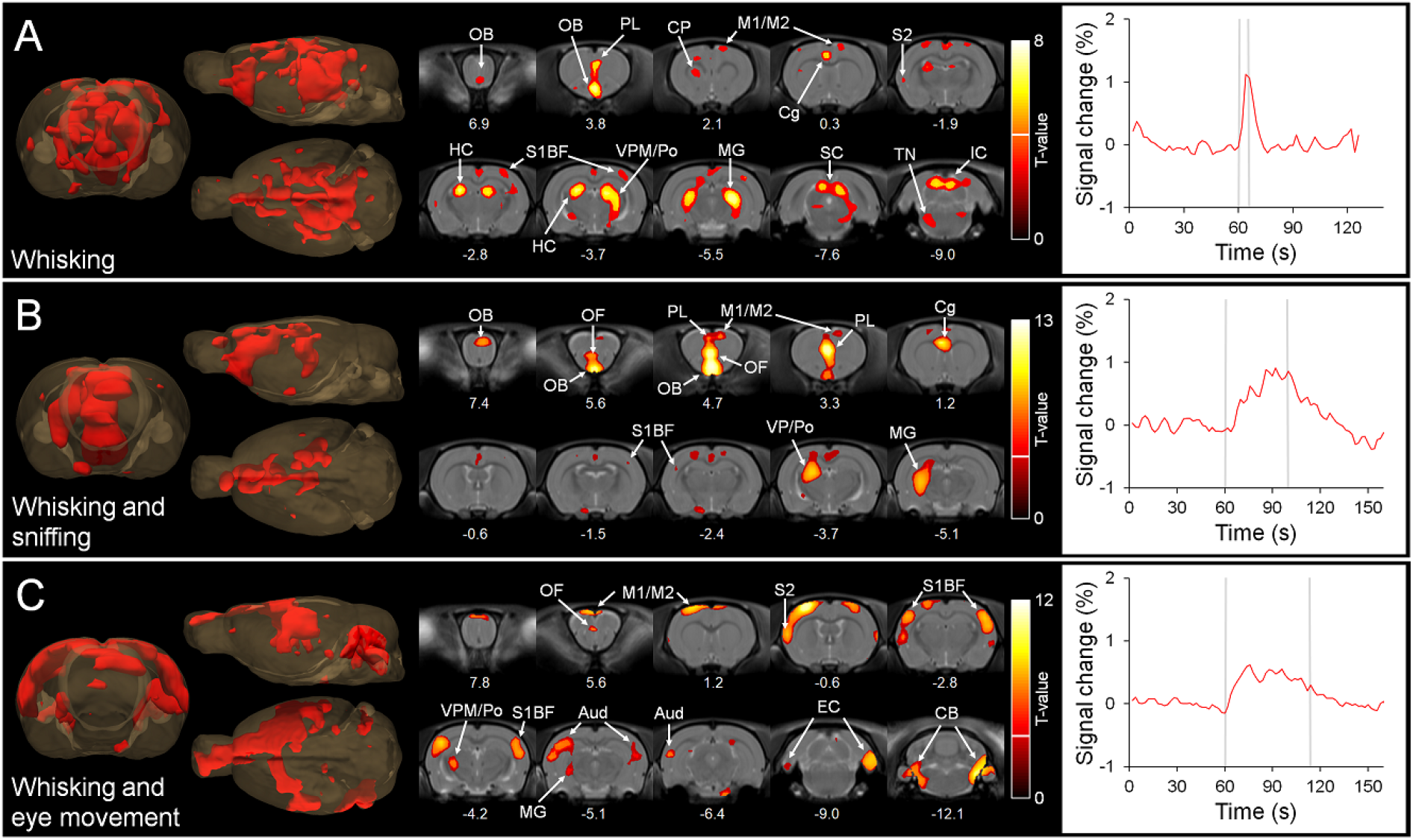
Representative statistical maps obtained from individual whisking-related behavioral events. In A, the rat was mainly whisking, with simultaneous activation of the cortico-thalamic barrel circuit and tectum (superior and inferior colliculus). In B, the rat was sniffing intensively, but also whisking, showing simultaneous activations in olfactory and barrel systems. In C, the rat was whisking in addition to eye movement, showing simultaneous activations in barrel and auditory cortices but not in the olfactory system. Time series represent average across all significant voxels at p < 0.001 (uncorrected). Gray bars in the time series indicate the window for the behavioral state observed in the video. Two statistical thresholds (p < 0.05 corrected with threshold indicated by the white line in each color bar; and p < 0.001 uncorrected corresponding to all colored voxels) were applied. Statistical maps are overlaid on SIGMA brain template, and numbers below slices indicate distance from bregma (mm). The 3D illustrations show views from the front, left side, and top of the brain. Aud, auditory cortex; BF, barrel field region; Cb, cerebellum; Cg, cingulate cortex; CP, caudate putamen; EC, entorhinal cortex; HC, hippocampus; IC, inferior colliculus; M1, primary motor cortex; M2, secondary motor cortex; MG, medial geniculate nucleus; OB, olfactory bulb; OF, orbitofrontal cortex; PL, prelimbic cortex; Po, posterior thalamic nuclei; S1, primary somatosensory cortex; S2, secondary somatosensory cortex; SC, superior colliculus; TN, trigeminal nuclei; VP; ventral posterior thalamus.

To extend the single behavioral event analyses further, we studied the brain activation patterns during unilateral limb use, startle behavior, and acute sustained change in alertness (Figure 6). During unilateral retreat attempt (Figure 6A), the rat tried to briefly twist itself free from the imaging holder by using mainly left hindlimb. Correspondingly, we observed the strongest activation in contralateral motor cortex, contralateral somatosensory hindlimb region, contralateral thalamus, and in ipsilateral cerebellum. Thus, the unique unilateral nature of the spontaneous behavior was well in line with the measured brain activity, further validating the methodology.

**Figure 6.**
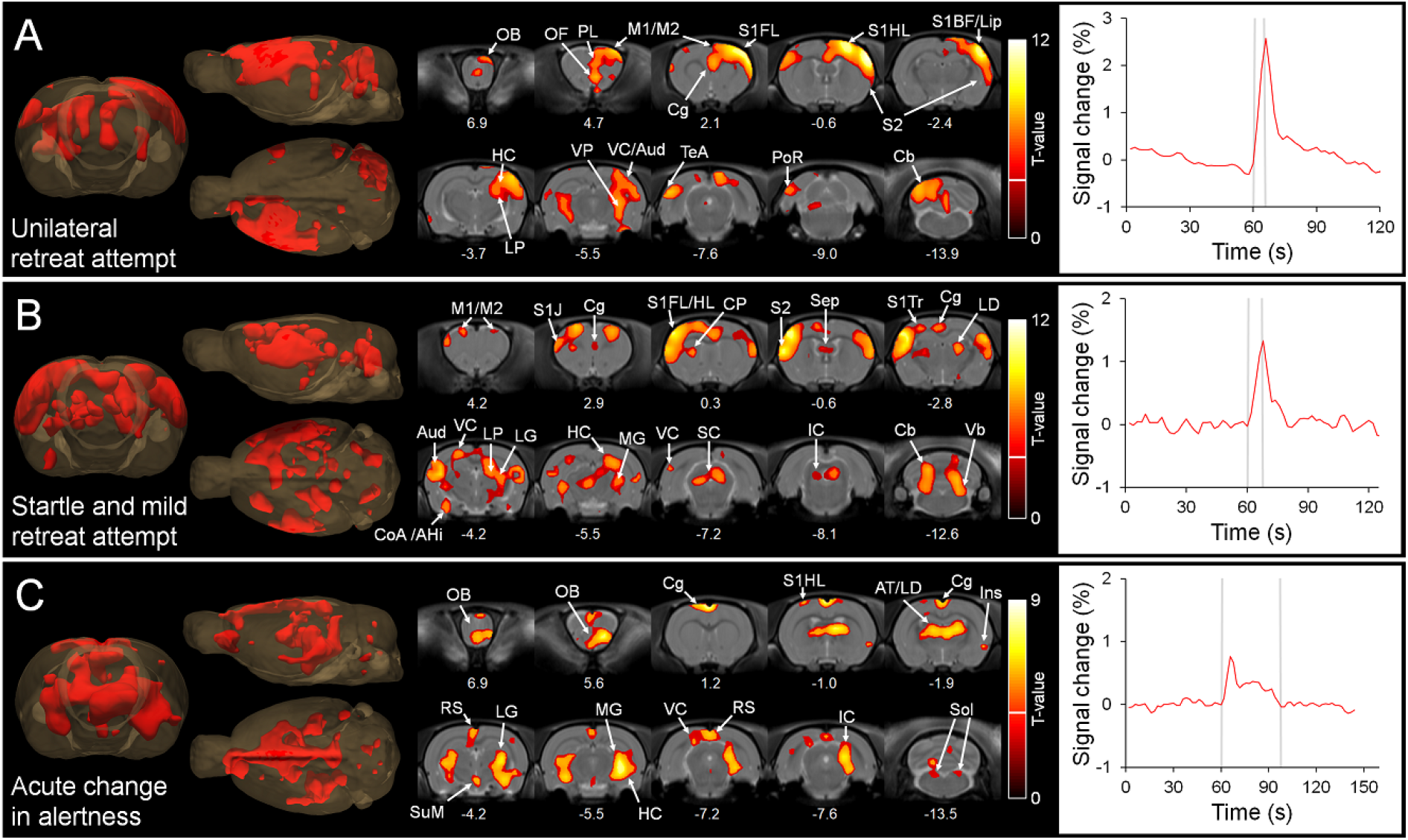
Representative statistical maps obtained from individual behavioral events (A-C). In A, the rat was mainly using its left hindlimb for applying force, with simultaneous activation of contralateral motor and sensory cortex and ipsilateral cerebellum. In B, the rat suddenly startled and took few steps backwards, with simultaneous activations in motor and sensory regions, vestibular nuclei, and cerebellum. In C, there was a sudden and relatively long change in general alertness level of the rat, including bruxing, eye blinking, twitches, and sniffing, with simultaneous activation in olfactory, visual, and motor systems. Time series represent average across all significant voxels at p < 0.001 (uncorrected). Gray bars in the time series indicate the window for the behavioral state observed in the video. Two statistical thresholds (p < 0.05 corrected with threshold indicated by the white line in each color bar; and p < 0.001 uncorrected corresponding to all colored voxels) were applied. Statistical maps are overlaid on SIGMA brain template, and numbers below slices indicate distance from bregma (mm). The 3D illustrations show views from the front, left side, and top of the brain. AT, anterior thalamic nuclei; Aud, auditory cortex; BF, barrel field region; Cb, cerebellum; Cg, cingulate cortex; CoA/AHi, cortical amygdaloid / amygdalohippocampal area; CP, caudate putamen; FL, forelimb region; HC, hippocampus; HL, hindlimb region; IC, inferior colliculus; J, jaw region; LD, laterodorsal thalamic nuclei; LG, lateral geniculate nucleus; Lip, lip region; LP, lateral posterior thalamic nuclei; M1, primary motor cortex; M2, secondary motor cortex; MG, medial geniculate nuclei; OB, olfactory bulb; OF, orbitofrontal cortex; PoR, postrhinal cortex; RS, retrosplenial cortex; S1, primary somatosensory cortex; S2, secondary somatosensory cortex; SC, superior colliculus; Sep, septum; Sol, nuclei of solitary tract; SuM, supramamillary nucleus; TeA, temporal association cortex; Tr, trunk region; VC, visual cortex; Vb, vestibular nuclei; VP, ventral posterior thalamic nuclei.

During the sudden startle (Figure 6B, Supplementary Video 3), we observed activation in the auditory (inferior colliculus, medial geniculate nuclei, and auditory cortex) and visual (superior colliculus, dorsal lateral geniculate nuclei, and visual cortex) pathways. We assume that a small environmental change led to startle, and signal changes in auditory and visual systems were related to the initiation of the behavior. Next, we observed signal changes in superior colliculus, anterior cingulate cortex, septum, and hippocampus that may be related to orientation because of the sudden environmental change. After the startle response, a mild retreat attempt was observed, which corresponds well to the observed activation of motor (motor cortex, caudate putamen, superior colliculus, and cerebellum) and sensory (primary and secondary somatosensory cortex, and lateral dorsal thalamic nuclei^42^) areas. Signal changes occurred also in vestibular nuclei that are important in body posture control^43^. Additionally, startle-related activation was observed in posteromedial amygdala, covering cortical amygdala and amygdalohippocampal area. Since the cortical amygdala has been associated with innate odor-driven avoidance or approach^44^, and no activation of the olfactory system was observed, a more likely signal source is the amygdalohippocampal area that was recently found to be related to aggressive behavior^45^. These observations indicate that the activation of multiple systems during even a single, short, and diverse behavioral event can be detected with MB-SWIFT fMRI.

In the last example, we noticed a sustained (30-40 s) elevation in general alertness in one of the rats, during which sniffing, twitching, bruxing, and moving/blinking eyes occurred, accompanied with an average increase in respiratory rate of 10 bpm. The fMRI signal changes (Figure 6C) were correspondingly observed in the olfactory bulb, thalamus (anterior, laterodorsal, and lateral and medial geniculate nuclei), inferior colliculus, and throughout the cingulate cortex (cingulate and retrosplenial cortex), consistent with attention towards potential olfactory, visual, or auditory stimuli. The lateral thalamic activation partly overlapped with the hippocampus. The involvement of the latter is supported by specific activation of the supramamillary hypothalamic nucleus, a major generator of hippocampal theta rhythm associated with active explorative state^46^. Additionally, signal increases in the insula and solitary tract nuclei may be related to sustained changes in autonomic functions such as breathing rate^47^. Again, the observed behavior corresponded well to the fMRI signal changes during a single event, and we did not observe global hemodynamic signal changes that could be related to the small acute increase in breathing.

Taken together, the results above emphasize the sensitivity of the novel methodology, as activation of relevant areas, including motor, somatosensory, auditory, visual, emotional, and autonomic control regions, were detected behavior-dependently during single events. Importantly, the majority of these activations were visible with the standard statistical threshold (p < 0.05, corrected).

## Discussion

Comprehending the underlying processes of normal and abnormal behavior is a challenging task that provides the basis for the development of treatments for mental diseases. To address such challenge, novel research approaches and concepts are commonly developed and tested in preclinical environments. In this work, our results indicate that the novel MB-SWIFT fMRI allowed, for the first time, the acquisition of anatomically and functionally relevant high-quality wholebrain data, including the hindbrain, in head-fixed spontaneously behaving rats. This could not be achieved with traditional EPI-based fMRI as it is highly sensitive to dynamic magnetic field changes^17,48^, making even breathing motion a factor that distorts images^49,50^. The minimal motion correction parameters, the functional data quality, and the analyses without brain masks all indicated that body movement was not a significant confounding factor with the MB-SWIFT fMRI approach. Furthermore, minor to no impact on image quality occurred because zero echo time and large bandwidth sequences are insensitive to susceptibility-induced image distortions^16^ originating from surgical manipulations, such as the head implant with acrylic cement and anchoring screws in this work. However, this is not the case with traditional EPI^17^, which may be one of the reasons why alternative and less robust head-fixation strategies are applied in the majority of the restrained awake rodent studies, which can subsequently deteriorate the data quality through increased head movement.

Although imaging of restrained rodents^14,20,51^ has limitations, the recent studies in restrained rats^17,25,30^ allowed us to validate the functional parcellation obtained in head-fixed rats. Importantly, similar functional network structure as observed here (Figures 2 and 3) and in previous work^25,30^, has not been reported in anesthetized rats, emphasizing the importance of awake imaging in functional studies. This is a serious concern because a majority of preclinical fMRI data have been acquired under anesthesia, which dramatically hinders the general interpretation and comparability of results^52^. While our functional parcellation and hierarchical clustering results were in good overall agreement with previous restrained awake studies^25,30^, emphasizing good reproducibility of awake rat imaging, a key difference was that we did not observe either static or spontaneous activity in the amygdala proper. The reason for this remains unclear, as one would expect stress or fear-related processing during the fMRI, and signal changes were observed in many related brain areas. We do not consider sensitivity as the restricting factor, as we detected signal changes in deep structures, such as in the olfactory tubercle and piriform cortex during sniffing. Furthermore, we did observe startle-related signal increases in amygdalohippocampal area, which abuts the amygdala proper. Therefore, the issue may be related to spatial resolution or the localization of the hemodynamic response involving larger vessels in close proximity to the activated region.

To the best of our knowledge, all behavior-related findings here are novel as there are no existing tools to study whole-brain activity in head-fixed rodents during spontaneous behavior. The group-level behavioral results suggest that 1) the observed spontaneous behavior corresponded to whole-brain activation patterns, 2) activation of multiple systems, such as sensory, decisionmaking and motor control, could be observed simultaneously, 3) the methodology is sensitive to frequently occurring sub-second events with < 1% signal changes (Figure 4C and Supplementary Video 1), and 4) distinct brain activation patterns with similar behavior could be distinguished (spontaneous vs. fruit odor-induced sniffing). Based on the anatomical localization of the fMRI signal changes and prior knowledge related to neuroanatomy, we conclude that the obtained activation maps were behaviorally relevant and likely reflecting the diverse behavior-related brain activity.

There are, however, few factors to be considered in our group-level behavioral analyses. First, the analysis of spontaneous behavior, during which the exact stimulus and its parameters remain unknown, is uncommon in fMRI. These unknown parameters include the cause, onset, and length of the behavior-related event in the brain. In the present work, the parameters used in analyses were based on visual observations, and the obtained activation maps suggested potential driving factors for the behavior in, e.g., sensory and emotional processing brain regions. While this might be accurate enough for simple approaches, complementary techniques, such as local field potential recordings, are required for more accurate spatio-temporal evaluations of the underlying parameters of spontaneous behavior. Fortunately, electrical recordings are fully compatible with our head-fixed approach, as shown here with the representative simultaneous EEG-fMRI measurements (Supplementary Figure 1). Second, the behavioral events are never identical, which induces variability in the group-level analyses. For example, the initial cause or its length leading to a retreat attempt from the imaging holder may vary greatly across the rats. Therefore, the group-level results mainly reflect the common brain activity patterns across subjects, for example in motor areas, and may leave activation in certain limbic or sensory regions undetected.

The best way to handle behavioral variability is to be able to study single events in individual animals, but this is challenging as statistical power in fMRI typically relies on repetitions and group-level data. However, we observed well-localized and relevant activation patterns during single behavioral events with the MB-SWIFT fMRI in head-fixed rats, suggesting excellent sensitivity of our approach. This is in good agreement with our previous work, where MB-SWIFT showed up to 6-fold higher functional signal-to-noise ratio compared to spin-echo EPI^16,18^. As the percentage signal changes in MB-SWIFT fMRI appear to be moderate (typically 0.5-1.0%), the high functional contrast likely originates from the very stable baseline. Importantly, the individual activation maps were able to demonstrate very specific aspects of individual behavioral events. For example, during the startle behavior when there was no sniffing, there was no activation in olfactory regions either. Furthermore, during acute alertness when no whisking occurred, there were no barrel circuit activation. However, we recognize that the sensitivity may not be high enough for robust detection of, e.g., all thalamic nuclei during each event, but at minimum the individual-level analysis may help to classify the data into proper subcategories if there are uncertainties related to the visually observed behavior.

While we recognize that studies in head-fixed animals have limitations related to the restricted native behavior and time-consuming training, there are several advantages compared to studies in freely running animals^19^, making the head-fixed approach relevant in behavioral neuroscience. Most importantly, head-fixed animals provide foundation for precise spatio-temporal studies of brain activity due to better control over both imaging and behavior, with minimal confounding effect of head movement. Additionally, studies do not require the use of anesthesia, which heavily modulates brain networks^52^, and can be performed without strict body restraint, which is a powerful stress-inducing factor in rodents^22,23^. Head-fixed condition allows numerous study designs related to, e.g., conditioned movement, Go/NoGo tasks, detection, discrimination, impulsivity, motivation, and reward system^19^. Even head-fixed exploration in virtual environment is possible^19,53^, which is one of the future challenges to be implemented with simultaneous fMRI. Nevertheless, we recognize that even a mild stress in head-fixed animals can modulate or prevent normal behavior^19^, and thus our habituation protocol requires further optimization to achieve as stress-free conditions as possible. This is likely achieved by simply extending the training period^19^.

Currently, there are no methods allowing the behavioral studies of brain function with similar scale and freedom as shown here. fUS is a modern approach that covers brain partially and has superior temporal and spatial resolution over fMRI^8,10^, making it an attractive tool for many applications in neuroscience. However, the biggest disadvantages of fUS are yet related to poor translatability due to limited bone penetration, making the approach currently compatible only with mice, human infants, or rats with a thinned skull. Importantly, the MB-SWIFT can be broadly implemented into existing preclinical and clinical scanners. The main requirement for the implementation is a correct-sized transmit-receive coil, as the 3D sequence has no slice selection and thus all excited signal needs to be covered inside the field of view to minimize artefacts originating from non-encoded signal^15^. The initial specific absorption rate calculations indicated that the absorbed energy by tissue during MB-SWIFT is acceptable for human use^16,17^. Moreover, conventional SWIFT has already been successfully applied in human fMRI with visual stimulation, where the robust and well-localized signal changes were in the same range as with gradient-echo EPI^54^. As the same limitations of traditional EPI fMRI, loud acoustic noise and movement-sensitivity, apply to both animals and humans, we anticipate that the implementation of MB-SWIFT, or other similar zero echo time large bandwidth sequence, into clinical use could have immediate positive impact on fMRI study designs and data quality. Subjects could perform more complex tasks that require body and limb movement, such as active conversation, eating, motoric coordination, or physical exercises. Moreover, as MB-SWIFT does not require highly homogenous main magnetic field due to its high bandwidth, completely new scanner designs could be achieved to enable a novel repertoire of whole-brain behavioral studies in humans. In theory, new scanners could be smaller doughnut-shaped devices surrounding the head and keeping it still, while the subject could sit, stand, have social interaction, or even walk on a roller mat and explore a virtual environment. Development towards such instruments is already ongoing^55^.

We conclude that novel zero echo time and large bandwidth fMRI applications, such as MB-SWIFT fMRI, can provide significant improvements into the currently inadequate whole-brain methodologies in behavioral research. These improvements can widen the available options in behavioral whole-brain research in rats, but also likely in humans. As EPI has been the choice for fMRI over almost three decades, we propose a shift towards more convenient, artefact-free, quiet, and stable MRI sequences with less restrictions in behavioral studies to facilitate the understanding of underlying mechanisms of behavior in the brain.

## Methods

### Animal Procedures and Experimental Timeline

Schematic presentation of the experimental timeline is shown in Supplementary Figure 1. All animal procedures were approved by the Finnish Animal Experiment Board and conducted in accordance with the European Commission Directive 2010/63/EU guidelines. Adult (448 ± 56 g at the onset of habituation) male rats (n = 10; Hsd:Sprague Dawley^®^ SD^®^; Envigo RMS B.V., Horst, Netherlands) allocated for the experiments were individually-housed and maintained on a 12/12 h light-dark cycle at 22±2 °C with 50%-60% humidity. Food and water were available ad libitum.

### Head Implant surgery

To allow firm head-fixation at multiple habituation and fMRI time points, a chronic implant compatible with a 22-mm diameter MRI transmit-receive single loop coil was attached on the skull of the rats. The protocol was similar to that described earlier^17^. Briefly, the rat was anesthetized with isoflurane (Attane vet 1000 mg/g, Piramal Healthcare UK Limited, Northumberland, UK; 5% induction and 2% maintenance) in 30/70% O_2_/N_2_ carrier gas. The scalp was removed from top of the skull, and small anchoring screws made of brass were inserted on top of both olfactory bulb and cerebellum. Additionally, small holes not penetrating the bone were drilled throughout the skull to facilitate further bone cement adhesion (Palacos R + G, Heraeus Medical, Hanau, Germany). A layer of dental cement (Selectraplus, DeguDent GmbH, Hanau, Germany) was applied on top of the bone cement. Two plastic pins (2 mm diameter) covered with heat-shrinkable tubes were fixed on top of the dental cement layer, and dental cement was molded over the pins. Subsequently, the pins were removed, leaving two holes for head-fixation. For two days twice a day, buprenorphine (0.03 mg/kg s.c.; Temgesic, Indivior Europe Ltd, Dublin, Ireland) and saline (10 ml/kg/day s.c.) were given for analgesia and rehydration, respectively. Rats recovered at least 1 week prior to the habituation for fMRI.

For one rat, EEG electrodes were implanted for simultaneous EEG-fMRI to follow the transition from anesthetized to awake state. Procedure followed our earlier report^56^. For the recording electrodes, both tips of insulated silver wire (0.27 mm diameter with coating, AG549511, Advent Research Materials, United Kingdom) were exposed for 2 mm. The connector end of the wire was soldered to a socket connector (E363/0, Plastics One Inc., Roanoke, VA), and the recording end was bent to form an open loop with 1 mm diameter. Brass screw electrodes were used as a reference and ground. The head-fixation implantation was similar as described above, except holes for the electrodes were drilled through the skull yet leaving dura intact. Subsequently, three measuring electrodes were placed over cortex (AP: +3, ML: +0.5; AP: +2, ML: +2.5; AP: −2, ML: +2.5 mm) and reference and ground electrodes were placed over cerebellum. The connector ends of electrodes were inserted into a plastic pedestal (MS363, Plastics One Inc.), which was positioned between the head-fixation pin holes and attached with dental cement to the implant.

### Habituation Protocol for Head-fixed awake Imaging

The schematic presentation of the progressive 9-day habituation protocol is shown in Supplementary Figure 1, which represents the gradual implementation of procedures and lengthening of habituation sessions. Rats were habituated individually 1-3 rats per day and were allowed to familiarize themselves with the handling person during the week preceding the actual protocol. Habituation was done in a home-made wooden box (90 x 40 x 30 cm^3^), equipped with the imaging holder, a loudspeaker (BG-17, Visaton, Haan, Germany), and an infrared web camera. The recording and playback (Servo 200, Samson, Hicksville, NY, USA) of scanner noise was done as described previously^14,17^. For sound pressure level comparison between EPI and MB-SWIFT, the scanner noise was recorded 30 cm away from the bore as the peak sound pressure level during EPI inside the bore exceeded the dynamic range of the recording microphone. Light maintenance anesthesia, either isoflurane (1.3-1.8%) or sevoflurane (1.8-3.0%; Sevorane, ABBVIE Oy, Helsinki, Finland), was used while inserting ear plugs and harness, while taking blood samples, and while the rat was positioned to or removed from the MRI holder. The glass surface of MRI animal holder was heated with warm water circulation (Haake SC 100, Thermo Fischer Scientific, Loughborough, England) and made slightly slippery by applying a thin layer of skin oil (Ceridal Lipolotion, Stada Nordic, Helsinki, Finland).

During the first three habituation days, the rat was allowed to freely roam and familiarize itself with the habituation box, commercially available walking harness (TRIXIE Heimtierbedarf GmbH & Co. KG, Tarp, Germany), silicone ear plugs, MRI holder, and scanner noise. On the fourth day, the handling person kept head still from the implant for short time periods (≤ 30 s) and gave reward immediately after releasing the head. The reward, given also always before and after the habituation or imaging, was either chocolate cereal or 1% sucrose water based on which one the rat preferred.

From the fifth to ninth habituation day, the rat was placed to the MRI holder with a custom-made head-fixation system for gradually lengthening periods. The head implant was fixed with two glass fiber pins and tightened from both sides, and harness was adjusted to appropriate height with Velcro^®^ straps sewn to the harness.

To evaluate the corticosterone levels during the habituation, blood samples were obtained from lateral tail vein at time points indicated by Supplementary Figure 1. Samples were handled, prepared, and analyzed with a corticosterone assay kit (Corticosterone (Human, Rat, Mouse) ELISA, IBL International GmbH, Hamburg, Germany) as described earlier^14^. Baseline samples were taken immediately before the first habituation session, and the rest of the samples were taken 10-20 min after finishing the day-specific habituation or imaging.

### Magnetic Resonance Imaging

MRI parameters were similar as described earlier^17^. Briefly, a 9.4 T magnet equipped with a 21-cm inner diameter gradient set and interfaced with an Agilent DirectDRIVE console (Palo Alto, CA, USA) was used. A 22-mm custom-made surface loop coil was used for both signal transmission and reception. Anatomical images were obtained with a fast spin-echo sequence with the following parameters: 3000 ms repetition time, 12 ms echo spacing with 8 echoes, 48 ms effective echo time, 4 averages, 62.5 kHz bandwidth, 256^2^ matrix size, 4.0 x 4.0 mm^2^ field of view, and 25 slices with 1-mm thickness. fMRI data were acquired with a radial 3D MB-SWIFT sequence with the following parameters: 2000 spokes with 0.97 ms repetition time resulting in a temporal resolution of ~2 s per fMRI volume, 192/384 kHz excitation/acquisition bandwidths, 6° flip angle, and 64^3^ matrix size with 4.0 x 4.0 x 4.0 mm^3^ field of view, resulting in a 625 μm isotropic resolution.

Prior to MRI, the rat was prepared and positioned to the MRI holder as during habituation, except the MRI surface loop coil was placed around the head implant below the head-fixation pins and a pneumatic breathing sensor pad (Model 1025, Small Animal Instruments Inc., New York, NY, USA) was placed inside the harness to follow breathing rate. An MRI-compatible video camera with infrared light (12M-i, MRC Systems GmbH, Heidelberg, Germany) was placed front side of the rat to record behavior. Warm water circulation (Corio CD, Julabo, Seelbach, Germany) kept the holder glass surface at 40°C. The rat was kept under shallow anesthesia (similar to habituation, breathing rate 60-90 bpm) during shimming, pulse calibration, and anatomical imaging. Subsequently, anesthesia was ceased, and functional imaging was started when breathing reached 100 bpm or rat moved spontaneously. After finishing fMRI, anesthesia was gradually increased, rat was removed from the holder, returned to its cage, and rewarded.

The task-free “resting-state” measurements were conducted during the first and second imaging days and consisted of 450-750 fMRI volumes (15-25 min) each. The length was chosen based on how well the animal performed during the habituation. Two rats were imaged three times, while one rat was imaged only once. During the third imaging day, the rats were exposed to fruit odor (gas flowing through a bottle containing 0.5% amyl acetate liquid; Sigma-Aldrich, Helsinki, Finland) for 1 min (30 volumes), preceding a 6-min baseline (180 volumes). Imaging was continued for 5 min (150 volumes) after the odor challenge, making total imaging time of 12 min (360 volumes). Two rats lost head implant before the third imaging day, and thus odor-challenge data were acquired from total of 8 rats.

In simultaneous EEG-fMRI experiments, EEG was recorded with a BrainAmp MR system (Brain Products GmbH, Gilching, Germany) with 5 kHz sampling rate. Raw signal was pre-amplified tenfold (Multi Channel Systems, Reutlingen, Germany).

The results related to the controlled body position change experiments presented here are obtained by reanalyzing our previously published data, and the experimental protocol is described in the corresponding report^17^.

### Data Processing and Analysis

Behavioral video data were analyzed with EthoVision XT 14.0 (Noldus, Wageningen, The Netherlands). Researcher marked manually starting and ending points for each behavioral event. Same person analyzed all data within each category to minimize user-induced variability.

MRI data were processed and analyzed with in-house made scripts, MATLAB (R2011a and R2018b; Mathworks Inc., Natick, MA, USA), Python (version 3.6.9; https://www.python.org/downloads/), FSL (version 5.0; https://fsl.fmrib.ox.ac.uk/fsl/fslwiki), FreeSurfer (FreeView version 2.0; https://surfer.nmr.mgh.harvard.edu/), and Aedes (Version 1.0 rev 219; http://aedes.uef.fi). The statistical maps were overlaid on SIGMA^57^ rat brain template (https://www.nitrc.org/projects/sigma_template). The MB-SWIFT fMRI data reconstruction, motion correction, co-registration, and smoothing were done similarly as previously^17^. Briefly, the MB-SWIFT raw data were reconstructed with SWIFT package 2018 (https://www.cmrr.umn.edu/swift/index.php). From each dataset, the first two volumes were discarded as signal was reaching steady state. As the surface loop coil produced a spherical intensity gradient into the MB-SWIFT volumes, each volume with a mask covering the brain and nearby tissue was corrected by using Advanced Normalization Tools (ANTs, http://stnava.github.io/ANTs/) N4 bias field correction^58^. Subsequently, the N4-corrected volumes were motion corrected using rigid co-registration scheme on antsRegistration^59^. The rigid transformations obtained from this step were applied to motion correct the original MB-SWIFT volumes. The individual datasets were co-registered to a reference template by first co-registering the anatomical images using ANTs with a chain of rigid, affine, and non-linear^60^ transforms. Subsequently, the acquired transformations were applied to MB-SWIFT images. Finally, data were spatially smoothed (0.6 mm full-width at half-maximum).

To minimize the effect of motion on correlation analyses, three nuisance regressors for both translation and rotation were obtained from motion correction parameter files. Additionally, individual-level independent component analysis (ICA; FSL MELODIC, https://fsl.fmrib.ox.ac.uk/fsl/fslwiki/MELODIC) was used to manually detect and regress out independent components that were clearly motion-related artefacts, such as those located only at the edges and having high-amplitude spikes in time series. To further minimize influence of non-biological signal fluctuations, time series obtained from the head implant was regressed out from each voxel. Finally, the data were inspected manually for remaining movement-related artefacts. Volumes with visually observable blurring or remaining motion (> 0.1 voxels mass center) were discarded. To maintain direct temporal comparability between the behavioral video and fMRI data, the discarded fMRI volumes were replaced with time-wise linearly interpolated volumes.

To obtain functional parcellation of head-fixed rats, group-level ICA was performed for all task-free data. The analysis was performed in two parts. First, a brain mask covering the olfactory bulb and cerebrum was made, and the number of components was set to 40. This provided a comparable approach to previous awake rat brain low-dimensionality parcellation^25^, which was used as a reference while validating the ICA results. Second, a brain mask for hindbrain, including cerebellum and brainstem, was made and the number of components was set to 30. As there are no references related to the functional parcellation of hindbrain, the number of components was selected to be within range of a typical selection in the literature (20-40)^17^. To minimize the inclusion of non-neural sources into the results, components localized at the surfaces or with poorly defined anatomical localization were excluded from the subsequent analyses and results.

For the functional connectivity structure analysis, ICA components were thresholded with the Z-scores given by the MELODIC and used as ROIs in the partial correlation calculations. The original MB-SWIFT signal was band-pass filtered at 0.01–0.15 Hz, and motion correction parameters and brain volume global signal were used as nuisance regressors to minimize the residual effect of movement. Fisher z-transformation was applied to correlation coefficients prior to statistical testing and averaging. Finally, functional modules were obtained by using hierarchical clustering with linkage-function in Python scipy-module.

For the group-level behavioral fMRI analyses, the timings for behavioral events obtained from the videos were transformed to event vectors and used in the event-based fMRI analysis. As individuals were typically imaged more than once, and as specific events only occurred in a subset of the acquired time series, we used a two-level approach, where each co-registered MB-SWIFT fMRI time series of an individual were first concatenated into a single continuous time series. Thus, the concatenated time series consisted of 1-3 time series depending on the number of separate imaging time-points for an individual. All detected events were modelled using linear general model design and analyzed in the first level using FSL FEAT^61^. The intensity differences of the separate acquisitions were regressed out and auto-regression noise model (1^st^ order polynomial) was used. The first level analysis was followed by a group-level test, where null hypothesis was that the mean response does not differ from zero. We used FMRIB’s Local Analysis of Mixed Effects modelling^62^ and estimation with 1 + 2 option, cluster-defining threshold, and family-wise error rate multiple comparison correction. Those individual-level time series that did not contain a certain event were excluded from the group-level analysis of that specific event. Individual subject analyses were done with a block-design general linear model available in Aedes. Rat brain atlas^63^ was used to obtain anatomical localizations during interpretation of the results.

The EEG data were processed and inspected using MATLAB. The signal was denoised from gradient-switching artefacts by using template-based removal approach^64^, where the template shape was estimated by using the first ten fMRI volumes in EEG data. Subsequently, the template was aligned and regressed throughout the EEG data.

### Statistics

All group-level values are represented as mean ± standard deviation. Statistical tests were done with two-tailed paired t-tests, or with third-party data analysis packages that are listed in the beginning of data analysis section. The threshold for statistical significance was set to p < 0.05, unless stated otherwise in the figure legends, and corrections for multiple comparisons were performed with either family-wise error rate approach or Benjamini-Hochberg false discovery rate^65^ correction.

## Abbreviations

ANTs: advanced normalization tools
BOLD: blood oxygenation level dependent
bpm: breaths per minute
EEG: electroencephalography
EPI: echo planar imaging
fMRI: functional magnetic resonance imaging
fUS: functional ultrasound
ICA: independent component analysis
MB-SWIFT: Multi-Band SWeep Imaging with Fourier Transformation
PET: positron emission tomography
ROI: region-of-interest

## Data Availability

Data and codes are available upon reasonable request.

## Acknowledgements

This work was supported by the National Institutes of Health grants U01-NS103569-01 and P41-EB027061, and Jane & Aatos Erkko Foundation.

## Author contributions

J.P, P.S, T.P, H.L, and O.G designed the study. J.P, P.S, H.L, and T.P conducted the *in vivo* work. D.I, M.G, S.Mi, and S.Ma assisted in conceptualization and methodology. J.P, P.S, H.L, E.P, R.S, H.T, and O.G participated in data processing, analysis, and interpretation. J.P and O.G supervised the work and wrote the original draft. O.G, S.Mi, and S. Ma contributed to funding acquisition and resources. O.G. administered the project. All authors contributed to manuscript review and editing process.

## Materials & Correspondence

Prof. Olli Gröhn, A.I.V. Institute for Molecular Sciences, University of Eastern Finland, P.O. Box 1627, FI-70211, Kuopio, Finland. Phone number: +358 50 359 0963. E-mail address

## References

1. Janak PH, Tye KM. From circuits to behaviour in the amygdala. Nature. 2015;517(7534):284–292.

2. Hogeveen J, Salvi C, Grafman J. ‘Emotional intelligence’: Lessons from lesions. Trends Neurosci. 2016;39(10):694–705.

3. Gallivan JP, Chapman CS, Wolpert DM, Flanagan JR. Decision-making in sensorimotor control. Nat Rev Neurosci. 2018;19(9):519–534.

4. Ziv Y, Ghosh KK. Miniature microscopes for large-scale imaging of neuronal activity in freely behaving rodents. Curr Opin Neurobiol. 2015;32:141–147.

5. Hanks TD, Summerfield C. Perceptual decision making in rodents, monkeys, and humans. Neuron. 2017;93(1):15–31.

6. Kerr JN, Nimmerjahn A. Functional imaging in freely moving animals. Curr Opin Neurobiol. 2012;22(1):45–53.

7. Kyme AZ, Angelis GI, Eisenhuth J, et al. Open-field PET: Simultaneous brain functional imaging and behavioural response measurements in freely moving small animals. Neuroimage. 2019;188:92–101.

8. Urban A, Dussaux C, Martel G, Brunner C, Mace E, Montaldo G. Real-time imaging of brain activity in freely moving rats using functional ultrasound. Nat Methods. 2015;12(9):873–878.

9. Han Z, Chen W, Chen X, et al. Awake and behaving mouse fMRI during go/no-go task. Neuroimage. 2019;188:733–742.

10. Rabut C, Correia M, Finel V, et al. 4D functional ultrasound imaging of whole-brain activity in rodents. Nat Methods. 2019;16(10):994–997.

11. Ferrier J, Tiran E, Deffieux T, Tanter M, Lenkei Z. Functional imaging evidence for task-induced deactivation and disconnection of a major default mode network hub in the mouse brain. Proc Natl Acad Sci U S A. 2020;117(26):15270–15280.

12. Becerra L, Pendse G, Chang PC, Bishop J, Borsook D. Robust reproducible resting state networks in the awake rodent brain. PLoS One. 2011;6(10):e25701.

13. Liang Z, King J, Zhang N. Uncovering intrinsic connectional architecture of functional networks in awake rat brain. J Neurosci. 2011;31(10):3776–3783.

14. Stenroos P, Paasonen J, Salo RA, et al. Awake rat brain functional magnetic resonance imaging using standard radio frequency coils and a 3D printed restraint kit. Front Neurosci. 2018;12:548.

15. Idiyatullin D, Corum CA, Garwood M. Multi-band-SWIFT. J Magn Reson. 2015;251:19–25.

16. Lehto LJ, Idiyatullin D, Zhang J, et al. MB-SWIFT functional MRI during deep brain stimulation in rats. Neuroimage. 2017;159:443–448.

17. Paasonen J, Laakso H, Pirttimaki T, et al. Multi-band SWIFT enables quiet and artefact-free EEG-fMRI and awake fMRI studies in rat. Neuroimage. 2020;206:116338.

18. Laakso H, Lehto LJ, Paasonen J, et al. Spinal cord fMRI with MB-SWIFT for assessing epidural spinal cord stimulation in rats. Magn Reson Med. 2021;86(4):2137–2145.

19. Schwarz C, Hentschke H, Butovas S, et al. The head-fixed behaving rat--procedures and pitfalls. Somatosens Mot Res. 2010;27(4):131–148.

20. King JA, Garelick TS, Brevard ME, et al. Procedure for minimizing stress for fMRI studies in conscious rats. J Neurosci Methods. 2005;148(2):154–160.

21. Chang PC, Procissi D, Bao Q, Centeno MV, Baria A, Apkarian AV. Novel method for functional brain imaging in awake minimally restrained rats. J Neurophysiol. 2016;116(1):61–80.

22. Harris RB, Mitchell TD, Simpson J, Redmann SM, Jr, Youngblood BD, Ryan DH. Weight loss in rats exposed to repeated acute restraint stress is independent of energy or leptin status. Am J Physiol Regul Integr Comp Physiol. 2002;282(1):R77–88.

23. García-Iglesias BB, Mendoza-Garrido ME, Gutiérrez-Ospina G, Rangel-Barajas C, Noyola-Díaz M, Terrón JA. Sensitization of restraint-induced corticosterone secretion after chronic restraint in rats: Involvement of 5-HT_7_ receptors. Neuropharmacology. 2013;71:216–227.

24. Kabir MM, Beig MI, Baumert M, et al. Respiratory pattern in awake rats: Effects of motor activity and of alerting stimuli. Physiol Behav. 2010;101(1):22–31.

25. Ma Z, Perez P, Ma Z, et al. Functional atlas of the awake rat brain: A neuroimaging study of rat brain specialization and integration. Neuroimage. 2018;170:95–112.

26. Wagner MJ, Luo L. Neocortex-cerebellum circuits for cognitive processing. Trends Neurosci. 2020;43(1):42–54.

27. Buzsáki G, Moser EI. Memory, navigation and theta rhythm in the hippocampal-entorhinal system. Nat Neurosci. 2013;16(2):130–138.

28. Slotnick B. Animal cognition and the rat olfactory system. Trends Cogn Sci. 2001;5(5):216–222.

29. Tsai PJ, Keeley RJ, Carmack SA, et al. Converging structural and functional evidence for a rat salience network. Biol Psychiatry. 2020;88(11):867–878.

30. Liu Y, Perez PD, Ma Z, et al. An open database of resting-state fMRI in awake rats. Neuroimage. 2020;220:117094.

31. Klockgether T, Schwarz M, Turski L, Sontag KH. The rat ventromedial thalamic nucleus and motor control: Role of N-methyl-D-aspartate-mediated excitation, GABAergic inhibition, and muscarinic transmission. J Neurosci. 1986;6(6):1702–1711.

32. Izquierdo A. Functional heterogeneity within rat orbitofrontal cortex in reward learning and decision making. J Neurosci. 2017;37(44):10529–10540.

33. Brockett AT, Tennyson SS, deBettencourt CA, Gaye F, Roesch MR. Anterior cingulate cortex is necessary for adaptation of action plans. Proc Natl Acad Sci U S A. 2020;117(11):6196–6204.

34. Nyhuis TJ, Masini CV, Day HE, Campeau S. Evidence for the integration of stress-related signals by the rostral posterior hypothalamic nucleus in the regulation of acute and repeated stress-evoked hypothalamo-pituitary-adrenal response in rat. J Neurosci. 2016;36(3):795–805.

35. Byrd KE. Characterization of brux-like movements in the laboratory rat by optoelectronic mandibular tracking and electromyographic techniques. Arch Oral Biol. 1997;42(1):33–43.

36. Rogers-Carter MM, Varela JA, Gribbons KB, et al. Insular cortex mediates approach and avoidance responses to social affective stimuli. Nat Neurosci. 2018;21(3):404–414.

37. Deschênes M, Moore J, Kleinfeld D. Sniffing and whisking in rodents. Curr Opin Neurobiol. 2012;22(2):243–250.

38. Wesson DW, Carey RM, Verhagen JV, Wachowiak M. Rapid encoding and perception of novel odors in the rat. PLoS Biol. 2008;6(4):e82.

39. Petersen CC. The functional organization of the barrel cortex. Neuron. 2007;56(2):339–355.

40. Gharaei S, Arabzadeh E, Solomon SG. Integration of visual and whisker signals in rat superior colliculus. Sci Rep. 2018;8(1):16445-018-34661-8.

41. Driscoll ME, Tadi P. Neuroanatomy, inferior colliculus. In: StatPearls. Treasure Island (FL): StatPearls Publishing LLC; 2021. NBK554468 [bookaccession].

42. Bezdudnaya T, Keller A. Laterodorsal nucleus of the thalamus: A processor of somatosensory inputs. J Comp Neurol. 2008;507(6):1979–1989.

43. de Waele C, Mühlethaler M, Vidal PP. Neurochemistry of the central vestibular pathways. Brain Res Brain Res Rev. 1995;20(1):24–46.

44. Root CM, Denny CA, Hen R, Axel R. The participation of cortical amygdala in innate, odour-driven behaviour. Nature. 2014;515(7526):269–273.

45. Sato K, Hamasaki Y, Fukui K, et al. Amygdalohippocampal area neurons that project to the preoptic area mediate infant-directed attack in male mice. J Neurosci. 2020;40(20):3981–3994.

46. Kirk IJ. Frequency modulation of hippocampal theta by the supramammillary nucleus, and other hypothalamo-hippocampal interactions: Mechanisms and functional implications. Neurosci Biobehav Rev. 1998;22(2):291–302.

47. Alheid GF, Jiao W, McCrimmon DR. Caudal nuclei of the rat nucleus of the solitary tract differentially innervate respiratory compartments within the ventrolateral medulla. Neuroscience. 2011;190:207–227.

48. Jezzard P. Correction of geometric distortion in fMRI data. Neuroimage. 2012;62(2):648–651.

49. Kalthoff D, Seehafer JU, Po C, Wiedermann D, Hoehn M. Functional connectivity in the rat at 11.7T: Impact of physiological noise in resting state fMRI. Neuroimage. 2011;54(4):2828–2839.

50. Bianciardi M, van Gelderen P, Duyn JH. Investigation of BOLD fMRI resonance frequency shifts and quantitative susceptibility changes at 7 T. Hum Brain Mapp. 2014;35(5):2191–2205.

51. Ferris CF, Smerkers B, Kulkarni P, et al. Functional magnetic resonance imaging in awake animals. Rev Neurosci. 2011;22(6):665–674.

52. Paasonen J, Stenroos P, Salo RA, Kiviniemi V, Grohn O. Functional connectivity under six anesthesia protocols and the awake condition in rat brain. Neuroimage. 2018;172:9–20.

53. Chen G, King JA, Lu Y, Cacucci F, Burgess N. Spatial cell firing during virtual navigation of open arenas by head-restrained mice. Elife. 2018;7:10.7554/eLife.34789.

54. Mangia S, Chamberlain R, De Martino F, et al. Functional MRI with SWIFT. Proc. Intl. Soc. Mag. Reson. Med. 2012;20:326.

55. Garwood M, Mullen M, Kobayashi N, et al. A compact vertical 1.5T human head scanner with shoulders outside the bore and window for studying motor coordination. Proc. Intl. Soc. Mag. Reson. Med. 2020:1278.

56. Pirttimaki T, Salo RA, Shatillo A, et al. Implantable RF-coil with multiple electrodes for long-term EEG-fMRI monitoring in rodents. J Neurosci Methods. 2016;274:154–163.

57. Barrière DA, Magalhães R, Novais A, et al. The SIGMA rat brain templates and atlases for multimodal MRI data analysis and visualization. Nat Commun. 2019;10(1):5699-019-13575-7.

58. Tustison NJ, Avants BB, Cook PA, et al. N4ITK: Improved N3 bias correction. IEEE Trans Med Imaging. 2010;29(6):1310–1320.

59. Avants BB, Tustison NJ, Stauffer M, Song G, Wu B, Gee JC. The insight ToolKit image registration framework. Front Neuroinform. 2014;8:44.

60. Avants BB, Epstein CL, Grossman M, Gee JC. Symmetric diffeomorphic image registration with crosscorrelation: Evaluating automated labeling of elderly and neurodegenerative brain. Med Image Anal. 2008;12(1):26–41.

61. Woolrich MW, Ripley BD, Brady M, Smith SM. Temporal autocorrelation in univariate linear modeling of FMRI data. Neuroimage. 2001;14(6):1370–1386.

62. Woolrich MW, Behrens TE, Beckmann CF, Jenkinson M, Smith SM. Multilevel linear modelling for FMRI group analysis using bayesian inference. Neuroimage. 2004;21(4):1732–1747.

63. Paxinos G, Watson C. The rat brain in stereotaxic coordinates. 7th ed. Elsevier inc.; 2014.

64. Allen PJ, Polizzi G, Krakow K, Fish DR, Lemieux L. Identification of EEG events in the MR scanner: The problem of pulse artifact and a method for its subtraction. Neuroimage. 1998;8(3):229–239.

65. Benjamini Y, Hochberg Y. Controlling the false discovery rate - A practical and powerful approach to multiple testing. Journal of the Royal Statistical Society. Series B: Methodological. 1995 (57):289–300.

