## Supplementary Figures for "Whole-brain studies of spontaneous behavior in head-fixed rats enabled by zero echo time MB-SWIFT fMRI"

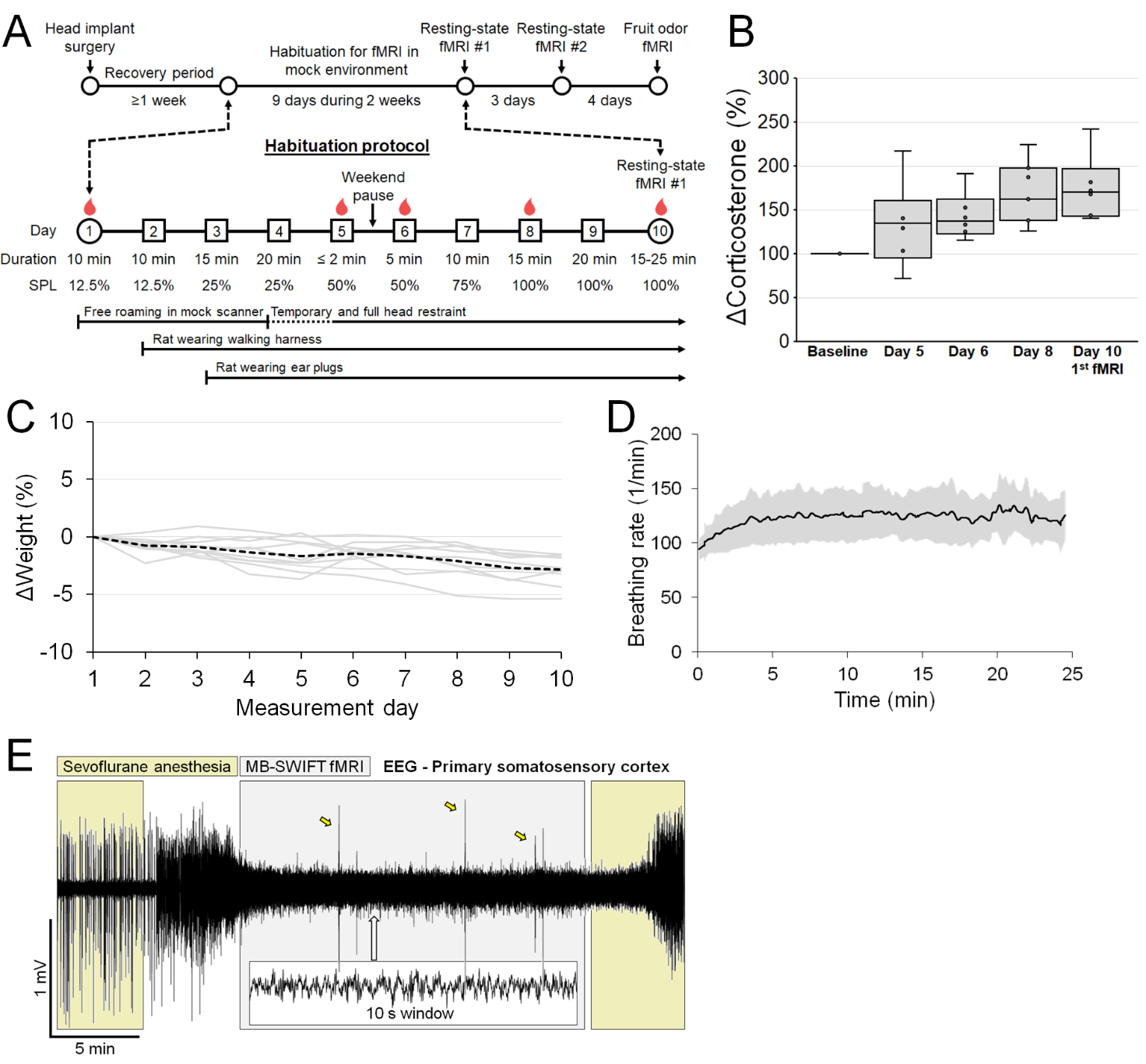
 **Supplementary Figure 1. The experimental timeline and habituation protocol (A), corticosterone levels (B) and weight (C) measured during the habituation, average breathing rate across fMRI measurements (D), and a representative electroencephalography (EEG) recording during simultaneous awake head-fixed MB-SWIFT fMRI (E).** In A, blood drop icons indicate corticosterone sampling days. Corticosterone levels increased (1.75-fold; p = 0.0012, paired t-test) and weight decreased (2.8%; p<0.001, paired t-test) mildly during the habituation protocol. However, there were no differences in corticosterone levels between day 8 and the first fMRI (p = 0.45, paired t-test). Average breathing rate (D) remained stable and similar to normal awake condition. An anesthesia-induced burst suppression state was clearly observed in cortical EEG before the awake imaging (E, first 5 min). After turning off the anesthesia, the brain shifted rapidly to awake state with theta-oscillations (4-8Hz). Only occasional movement-induced spiking artefacts were observed in the EEG data (E, yellow arrows). SPL, sound pressure level.

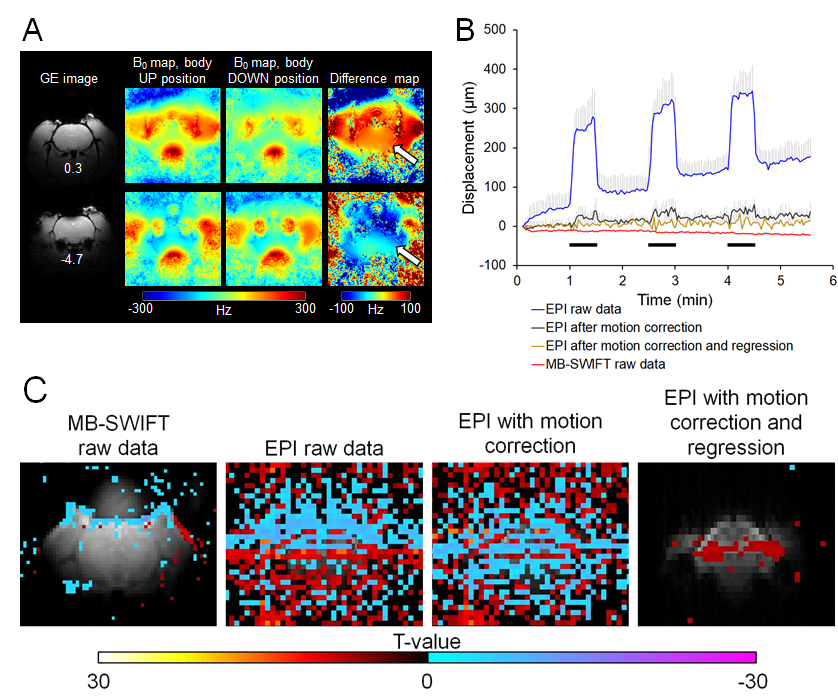
**Supplementary Figure 2. Effect of body position change on B_0_ field (A), center of mass displacement (B), and fMRI voxels (C).** The results presented here are re-analyses of previously published data, and methods are described in the corresponding report^17^. In controlled body position change experiments, the rear body of head-fixed and isoflurane-anesthetized rat was lifted 10-20 mm remotely during measurements, mimicking a body position change occurring during spontaneous behavior. The head did not move physically. White arrows in A indicate the position-related B_0_ field distortions in two representative slices in the rat brain. The black bars in B indicate the timing for controlled body position changes that greatly affected EPI data but had minimal effect on MB-SWIFT data (n = 6 in each). The center of mass displacement is plotted from the EPI phase-encoding direction (up-down). In C, voxels affected by body position change (block design with general linear model, p < 0.01, false discovery rate corrected) are shown in representative data sets. The raw MB-SWIFT data shows body position-related signal changes mainly in the skin, which is physically moving due to stretching. In contrast, EPI data suffers from significant signal changes inside the brain despite motion correction or motion correction following the regression of motion correction parameters prior to analysis.

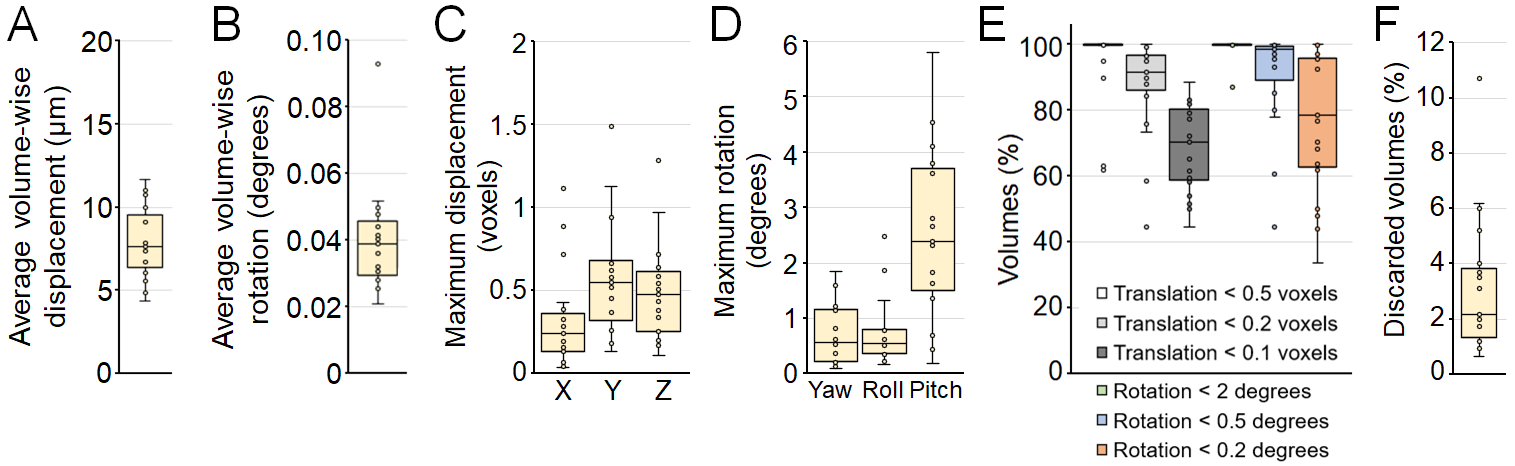
 **Supplementary Figure 3. Average volume-wise displacement (A) and rotation (B), maximum displacement (C) and rotation (D), and the amount of corrected (E) and discarded (F) volumes.**  Values in A and B indicate the translation and rotation compared to previous volume. Values in C-E are derived from motion correction parameters and are thus in relation to the original position and direction at the onset of imaging. Values in F are a result of manual inspection of the data after motion correction, indicating volumes that were not corrected successfully and were discarded. Each mark represents one data set (total n = 21). Marks outside whiskers are considered outliers.

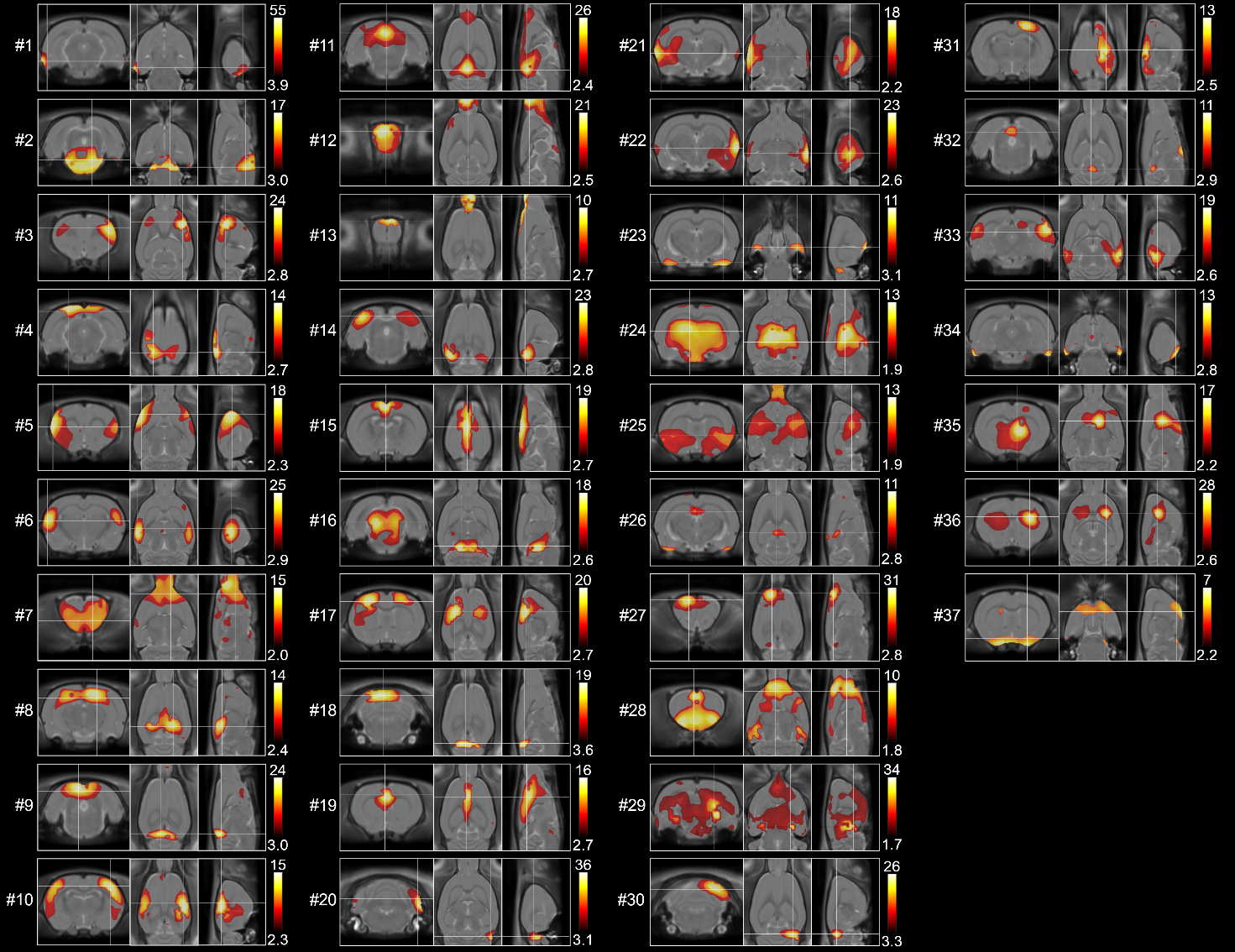
**Supplementary Figure 4. Independent components obtained from cerebrum and olfactory bulb with group-level ICA (n = 21).** Analysis was run with 40 components, and three components were excluded due to clear non-physiological origin. The numbering of components is the same as provided by the MELODIC, thus representing how much the component explains variance. The numbering does not match with the manuscript Figure 2, as some of the components were merged into the same subfigure for illustration purposes in Figure 2. Therefore, the number of figures does not match. ICA, independent component analysis.

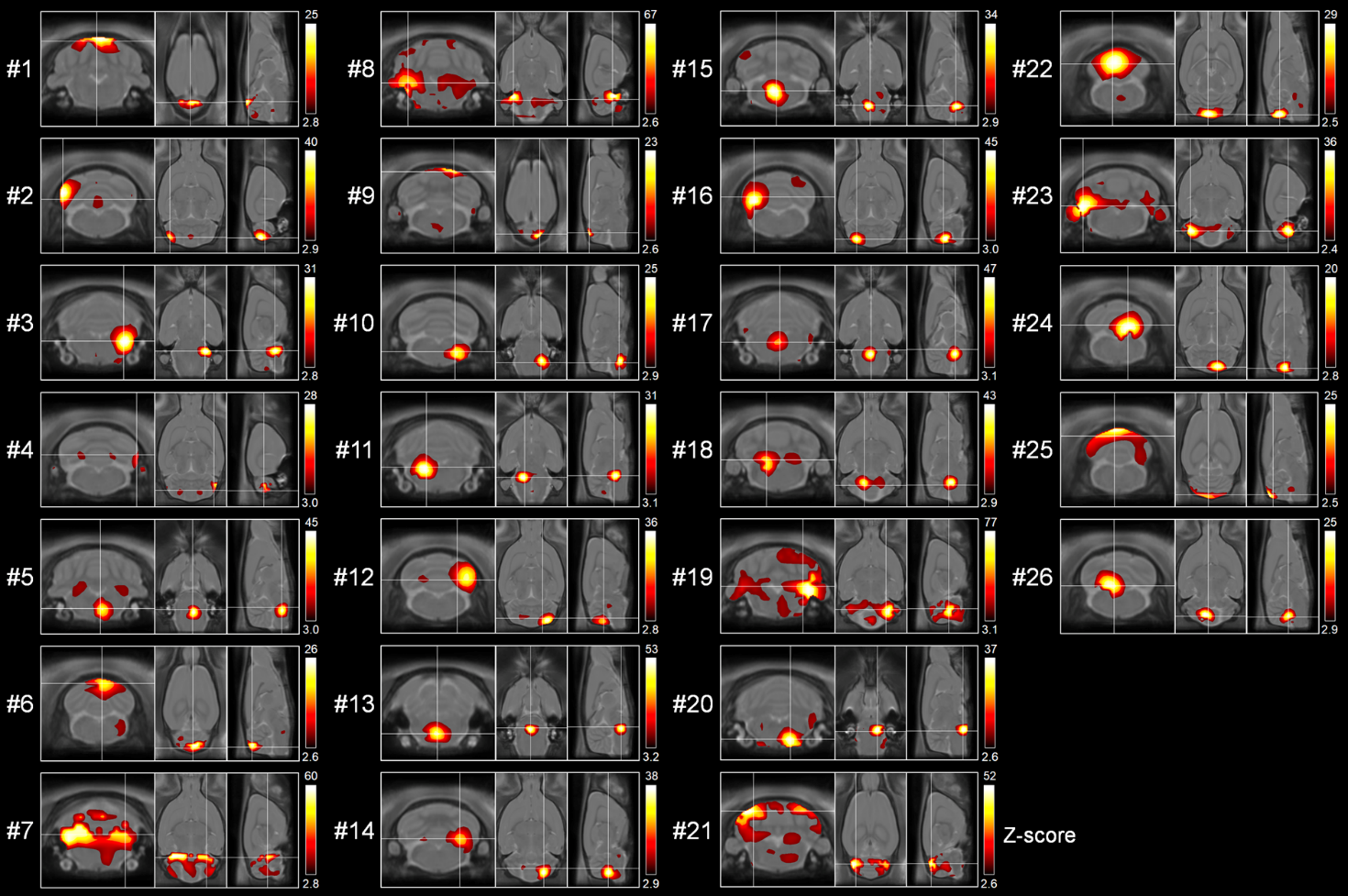
 **Supplementary Figure 5. Independent components obtained from hindbrain with group-level ICA (n = 21).** Analysis was run with 30 components, and four components were excluded due to clear non-physiological origin. The numbering of components is the same as provided by the MELODIC, thus representing how much the component explains variance. The numbering does not match with the manuscript Figure 2, as some of the components were merged into the same subfigure for illustration purpose in Figure 2. Therefore, the number of figures does not match. ICA, independent component analysis.

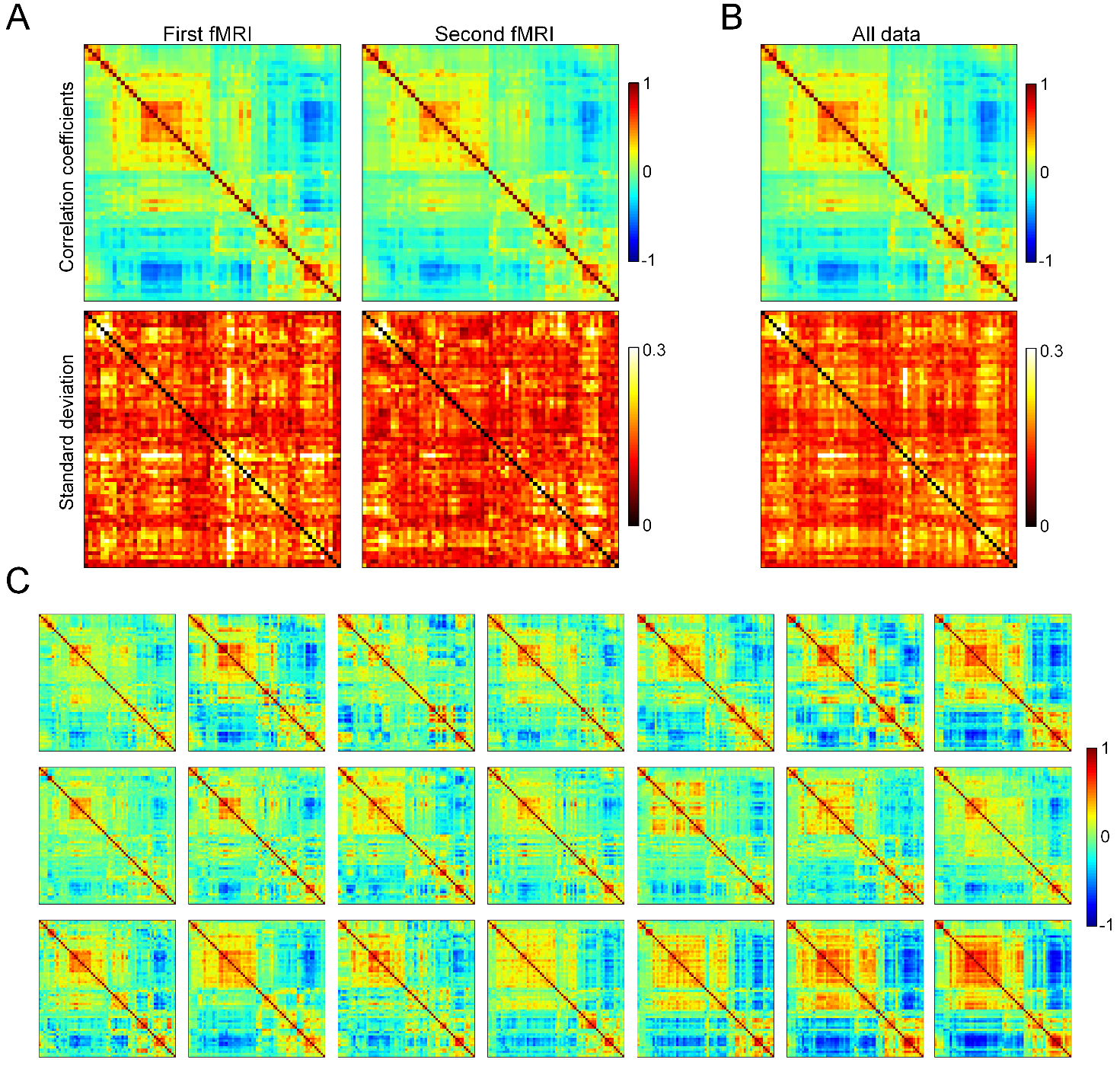
 **Supplementary Figure 6. Group- (A-B) and individual-level (C) partial correlation matrices.** In A, average and standard deviation matrices obtained during the first and second fMRI run are shown. In B, all individual-level data (shown in C) are pooled together. Regions-of-interest were derived from data-driven independent component analysis (Supplementary Figures 3 and 4) and organized by using hierarchical clustering.

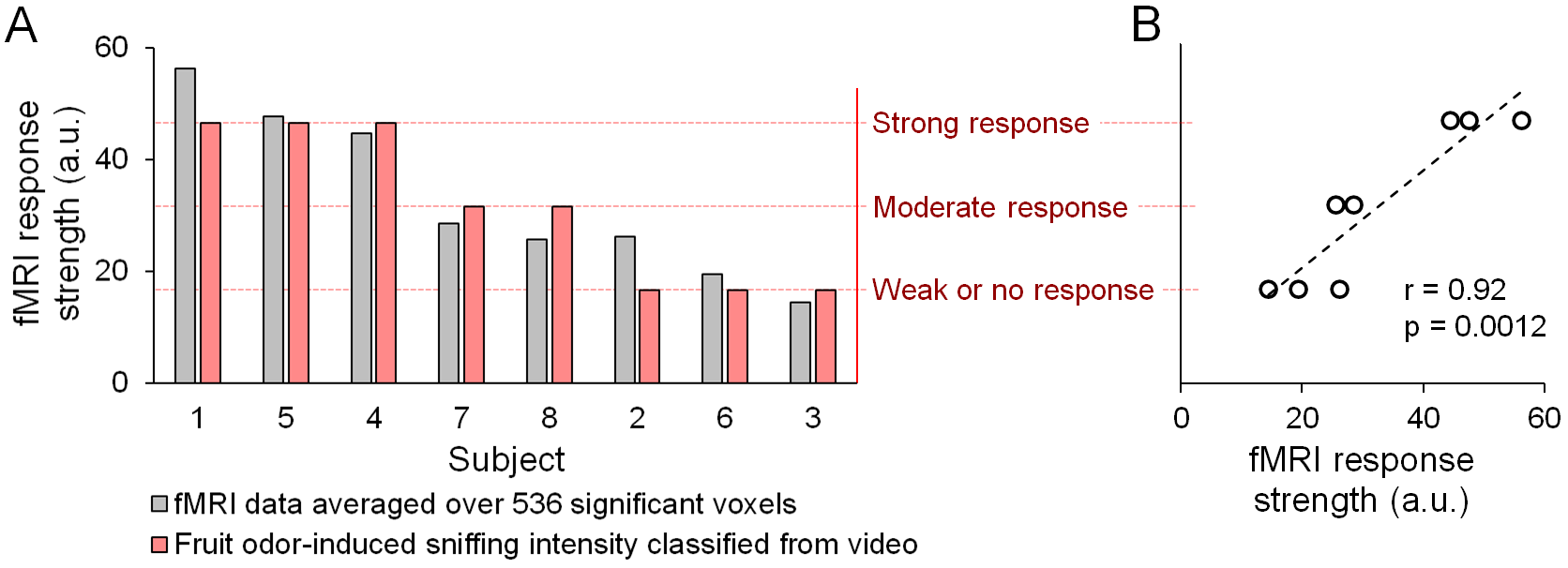
 **Supplementary Figure 7. fMRI response strength and classification of behavioral response to fruit odor (A), and their correlation (B).** Behavioral response was classified in one of three categories from the video and used as a regressor in fMRI analysis (FSL FEAT) to account for individual variation in the response strength. Classification and fMRI analyses were done by separate people that were not aware of the results.

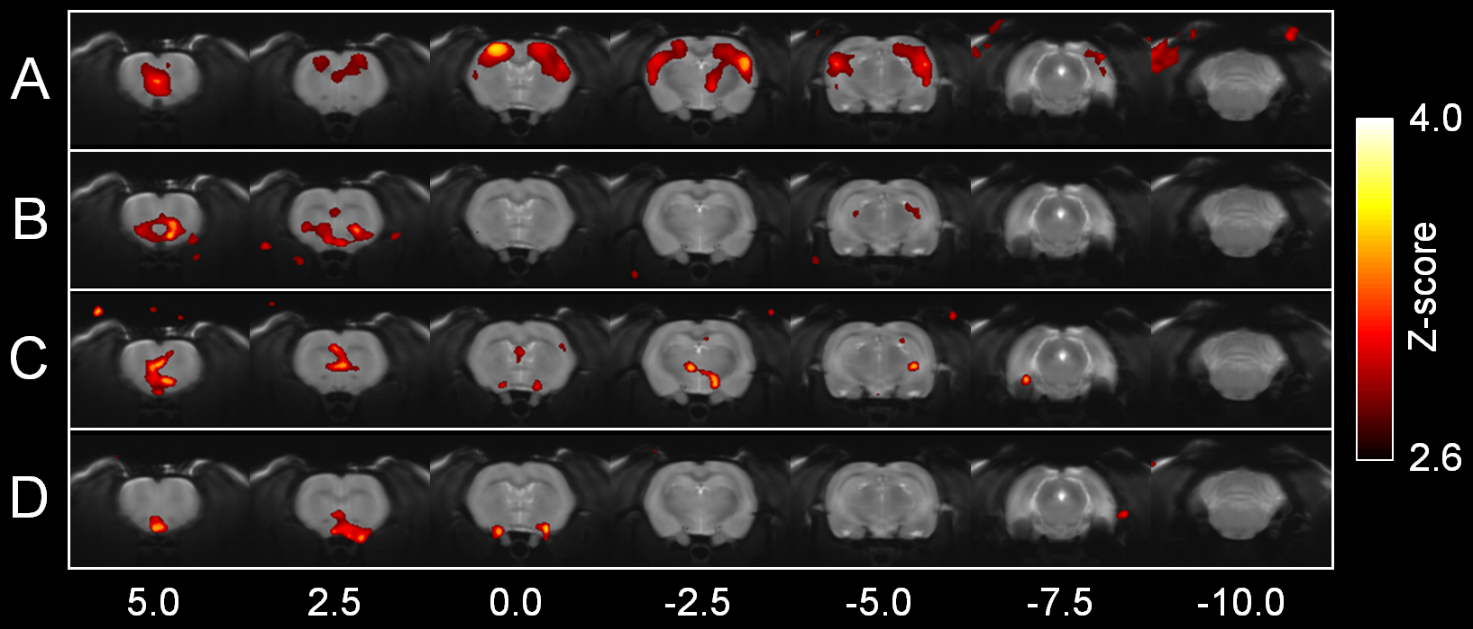
 **Supplementary Figure 8. Group-level spontaneous behavior-related statistical activation maps (A-D) without brain mask.** The groups are same as in Figure 4 in manuscript. In A, the rat tried to retreat. In B, the rat was bruxing. In C, the rat was sniffing, And in D, the rat was given fruit-odor. Only minor activation is observed outside the brain, mainly near the skin and ears that are typically moving while the rat is behaving. Statistical maps are overlaid on T_2_-weighted fast-spin echo images acquired prior to the MB-SWIFT fMRI scans. The slight mismatch of activation maps and the FSE images, particularly in the areas next to air cavities and skull, originates from the susceptibility-induced signal void in FSE image that is not present in the MB-SWIFT images. Numbers below slices indicate distance from bregma.
